# Molecular and cellular rhythms in excitatory and inhibitory neurons in the mouse prefrontal cortex

**DOI:** 10.1101/2024.07.05.601880

**Authors:** Jennifer N. Burns, Aaron K. Jenkins, RuoFei Yin, Wei Zong, Chelsea A. Vadnie, Lauren M. DePoy, Kaitlyn A Petersen, Mariya Tsyglakova, Madeline R. Scott, George C. Tseng, Yanhua H. Huang, Colleen A. McClung

## Abstract

Previous studies have shown that there are rhythms in gene expression in the mouse prefrontal cortex (PFC); however, the contribution of different cell types and potential variation by sex has not yet been determined. Of particular interest are excitatory pyramidal cells and inhibitory parvalbumin (PV) interneurons, as interactions between these cell types are essential for regulating the excitation/inhibition balance and controlling many of the cognitive functions regulated by the PFC. In this study, we identify cell-type specific rhythms in the translatome of PV and pyramidal cells in the mouse PFC and assess diurnal rhythms in PV cell electrophysiological properties. We find that while core molecular clock genes are conserved and synchronized between cell types, pyramidal cells have nearly twice as many rhythmic transcripts as PV cells (35% vs. 18%). Rhythmic transcripts in pyramidal cells also show a high degree of overlap between sexes, both in terms of which transcripts are rhythmic and in the biological processes associated with them. Conversely, in PV cells, rhythmic transcripts from males and females are largely distinct. Moreover, we find sex-specific effects of phase on action potential properties in PV cells that are eliminated by environmental circadian disruption. Together, this study demonstrates that rhythms in gene expression and electrophysiological properties in the mouse PFC vary by both cell type and sex. Moreover, the biological processes associated with these rhythmic transcripts may provide insight into the unique functions of rhythms in these cells, as well as their selective vulnerabilities to circadian disruption.

**Significance statement:** This is the first study to examine translatomic rhythms in the mouse PFC with cell-type specificity. We find that the core molecular clock cycles in phase across cell types, indicating that previously described daily oscillations in the cortical excitation/inhibition balance are not the consequence of a phase offset between PV and pyramidal cells. Nevertheless, rhythmic transcripts and their associated biological processes differ by both sex and cell type, suggesting that molecular rhythms may play a unique role in different cell types and between sexes. Therefore, our results, such as the enrichment of transcripts associated with mitochondrial function in PV cells from males, point towards possible cell and sex-specific mechanisms that could contribute to psychiatric and cognitive diseases upon rhythm disruption.

## Introduction

The PFC is a highly interconnected brain region that is involved in the integration of information, cognitive function, and emotional regulation (Miller, 2000; Fuster, 2001; Carlén, 2017). The predominant cell type in the PFC is glutamatergic pyramidal cells, which provide both local and long-range excitation and thus serve to carry the output of the PFC across the brain (Le Merre et al., 2021). Additionally, a variety of inhibitory interneurons modulate pyramidal cell function. Among these are parvalbumin (PV) expressing interneurons, which are also referred to as fast-spiking interneurons due to their ability to fire high frequency trains of non-adapting action potentials. These cells provide powerful inhibition to the soma and axon initial segment of pyramidal neurons and therefore act to control their output and regulate the overall excitation/inhibition (E/I) balance (Rudy et al., 2011; Hu et al., 2014; Ferguson and Gao, 2018). Notably, disruption of the PV-pyramidal cell microcircuit in the PFC in thought to underlie the cognitive symptoms of schizophrenia (Lewis et al., 2012; Gonzalez-Burgos et al., 2015; Chung et al., 2016, 2022; Smucny et al., 2022).

Circadian or diurnal rhythms are ∼24-hour rhythms that allow an organism to optimize the timing of physiological processes with the environment. The superchiasmatic nucleus (SCN) acts as the master pacemaker, synchronizing rhythms across the body. At the molecular level, these rhythms are orchestrated by a transcriptional-translational feedback loop that consists of the CLOCK and BMAL1 proteins, which dimerize and activate the transcription of *Period* (*Per*) and *Cryptochrome* (*Cry*) genes. In turn, the PER and CRY proteins dimerize and inhibit CLOCK and BMAL1 activity (Buhr and Takahashi, 2013). Beyond their role in the regulation of the core molecular clock, these proteins also promote the transcription of thousands of downstream clock-controlled genes, and in doing so, produce rhythms in the expression of many transcripts, including those associated with basic cellular functions (Oishi et al., 2003; Miller et al., 2007; Mure et al., 2018). Multiple studies have found that this rhythmicity is sexually dimorphic, with sex differences in the timing/amplitude of core clock genes and in broad transcriptomic rhythms (Lim et al., 2013; Chun et al., 2015; Logan et al., 2022; Talamanca et al., 2023). Moreover, in humans, an examination of rhythmic gene expression in the PFC found that rhythmic transcripts are associated with different functional pathways in males and females (Logan et al., 2022), suggesting that there may be biological processes with strong circadian rhythms in one sex and little daily fluctuation in the opposite sex.

Previous studies in the mouse SCN have found that rhythmic gene expression varies by cell type (Wen et al., 2020). These cell-type specific rhythms may drive the unique activity patterns of individual cell types, as rhythmic gene expression in the SCN is shifted in cell types known to have opposing rhythms in activity (Brancaccio et al., 2017; Wen et al., 2020). Differences in cell-type specific diurnal rhythms may also be present in the cortex, where daily shifts in the E/I balance may relate to rhythms in cognitive function (Schmidt et al., 2007; Chellappa et al., 2016; Bridi et al., 2020). Indeed, a recent study in the mouse PFC found that there is sex-specific diurnal variation in the electrophysiological properties of pyramidal neurons (Roberts and Karatsoreos, 2023), suggesting that the function of these cells may be dependent on time of day. The molecular basis of these rhythms has not been determined, as no studies of the PFC have examined cell-type specific rhythms in gene expression. In this study, we isolated ribosome associated transcripts from samples enriched in PV and pyramidal cells from the PFC of male and female mice across 24 hours to assess cell-type and sex-specific patterns in rhythmic gene expression. Additionally, we performed electrophysiological recordings from PV cells to assess whether there are similar diurnal changes in excitability and action potential characteristics as have been previously described in pyramidal cells (Roberts and Karatsoreos, 2023). We find that while rhythms in transcripts associated with the molecular clock are conserved and synchronized across cell types, broad translatomic rhythms differ by cell type and by sex, particularly in PV interneurons. Likewise, PV cells from female mice show diurnal rhythms in their electrophysiological properties that are absent after environmental circadian desynchronization (ECD).

## Methods

### Animals

To isolate actively translated transcripts enriched in PV and pyramidal cells respectively, *PV::cre* (Jax: 017320) and *CamKIIα*::*cre* (Jax: 005359)(Tsien et al., 1996) mice were crossed to Ribo-tag mice (Jax: 011029)(Sanz et al., 2009), which express a hemagglutinin (HA) tag on ribosomes in a cre dependent manner. Mice used for sequencing were group housed in a 12:12 light:dark (L:D) cycle (lights on 0700, Zeitgeber time (ZT) 0, lights off 1900 (ZT12) with access to food and water *ad libitum*. For PV cell electrophysiology experiments, G42 mice (Jax: 007677)(Chattopadhyaya et al., 2004) were bred to non-carrier G42 mice for recordings under a normal L:D cycle or to wild type mice on a mixed BALB/cJ x C57BL/6 background for ECD recordings. Upon weaning, mice to be used for recordings during the dark phase were transferred to a reverse light room (lights off 0700 (ZT12), lights on 1900 (ZT0), while animals to be used for light phase recordings remained under the normal L:D cycle. For ECD recordings, female mice were placed under a 10:10 L:D cycle (lights on ZT0, lights off ZT10) for a minimum of 3 weeks prior to recordings, as described in (Roberts and Karatsoreos, 2023). All mice used for these experiments were adults (∼13 weeks) and all experiments were performed in accordance with University of Pittsburgh Institutional Animal Care and Use Committee guidelines.

### Cell-type specific RNA isolation for rhythmicity studies

Male and female mice heterozygous for either *PV::cre* or *CamKIIα::cre* and carrying one copy of the RiboTag allele were sacrificed by cervical dislocation in 4 hour intervals across 24 hours (ZT 2, 6, 10, 14, 18, and 22); brains were subsequently removed and placed on dry ice. Serial sections were cut (150µm) on a cryostat (Leica Biosystems, Wetzlar, Germany) and tissue punches containing the anterior cingulate cortex and prelimbic region were taken. To isolate actively translated transcripts from PV and pyramidal cells, a co-immunoprecipitation protocol was used as previously described in (Chandra et al., 2015; Becker-Krail et al., 2022; DePoy et al., 2024), with minor modifications. Briefly, tissue was homogenized and incubated overnight with an anti-hemagglutinin A (HA) antibody (Abcam, Cambridge, UK). HA-tagged ribosomes, conjugated to an anti-HA antibody, were then isolated using Dynabeads (Invitrogen, Waltham, MA, USA). RNA was extracted and purified using an EZNA RNA Isolation Kit (Omega Bio-Tek Total RNA Miniprep kit) with ENZA RNase-Free DNAase (Omega Bio-Tek, Norcross, GA, USA).

### Sequencing

At the University of Pittsburgh Health Sciences Sequencing Core (Pittsburgh, PA, USA), isolated RNA was assessed for concentration (HSRNA Qubit, Thermo Fisher Scientific, Waltham, MA, USA) and integrity (Agilent RNA 6000 Pico kit; Agilent Technologies, Santa Clara, CA, USA). The average RNA integrity number (RIN) for samples from both cell types was high (>9). cDNA was then synthesized using the SMART-seq HT kit (Takara Bio, Inc, Kusatsu, Japan); one sample (PV M ZT18) was removed due to inability to generate cDNA. Library prep was performed using Nextera Flex (Illumina, San Diego, CA, USA) and samples were sequenced (2×101bp, 40 million reads/sample) using a Nova-Seq S4 (Illumina) at the UPMC Genome Center (Pittsburgh, PA, USA). Generated reads were evaluated for quality using FASTQC and were aligned to the mouse reference genome (*Mus musculus* Ensembl GRCm38) using HISAT2. HTSeq was used to convert reads to expression count data. To generate a list of transcripts to compare across cell types, transcripts were filtered across groups and transformed to log_2_CPM (counts per million) using the Bioconductor edgeR package. Only transcripts with a log_2_CPM>1 in >50% of the samples in at least one cell type were used for downstream analysis; Y chromosome linked transcripts were also excluded. This resulted in a background list of 13102 transcripts to be used for further analysis. Discrete clustering of cell types based on expression profile was performed using PCAtools (R software) to confirm that transcripts isolated from PV and pyramidal cells form segregated groups. Samples beyond 0 on the X axis and above 100 on the Y axis in the principal component analysis (PCA) were removed as outliers as their expression profile varied from other samples of the same cell type. This resulted in the exclusion of 4 PV cell samples (PV ZT2 F, PV ZT2 M, PV ZT6 M, PV ZT10 F) and one pyramidal cell sample (PYR ZT18 M), leaving 4-5 samples per timepoint per sex for downstream analyses.

### Experimental design and statistical analyses of RNA-sequencing data

Differential expression (DE) analysis was performed on count data using limma-voom by DESeq2 (Love et al., 2014). Transcripts were considered differentially expressed (enriched) with a log_2_fold change (log_2_FC) >0.58 and FDR corrected q-value <0.01. For rhythmicity analysis, the median of ratios normalization method in DESeq2 (Love et al., 2014) was used to normalize for sequencing depth and RNA composition, followed by log_2_ transformation. Transcript expression across time of day was fitted to a sinusoidal curve using a parametric cosinor model for rhythmicity analysis (Cornelissen, 2014). The R^2^ value was used as a measure of goodness of fit, with p-values calculated from the F-test. Rank-rank hypergeometric overlap (RRHO) was used as a threshold-free approach to assess overlap in rhythmic transcripts (Cahill et al., 2018). In this method, transcripts from two groups are ordered by significance (-log_10_p-value) along the X and Y axis, with the most significant transcripts in the bottom left corner. Overlap in the resulting heat map is indicated by a color gradient. Additionally, overlap in rhythmic transcripts between groups was compared using a p-value cutoff of p<0.0.5 and assessed using Fisher’s Exact Test and Odds Ratio. We considered transcripts that met the following stringent criteria to be uniquely rhythmic: (a) be significantly rhythmic (p<0.05) in one group, (b) have a significantly higher goodness of fit (R^2^) in the same group in which the transcript is rhythmic, and (c) not be significantly rhythmic in the other group (p>0.05).

To examine the biological processes associated with rhythmic transcripts, we used Ingenuity Pathway Analysis (IPA) (Qiagen, Hilden, Germany). For each analysis, rhythmic transcripts (p<0.05) were analyzed against a user supplied background list of transcripts. To compare enriched (p<0.05, log_10_p-value>1.3) pathways across groups, the top enriched pathways in each group were assessed for enrichment in the opposite group. For analysis of transcripts by phase, rhythmic transcripts (p<0.05) that peaked between ZT0-12 were defined as light phase, while rhythmic transcripts that peaked between ZT12-24 were defined as dark phase. For all analyses, pathways containing fewer than 15 genes were excluded. The top 10 enriched pathways are shown for each analysis except in the analysis of PV cells in the light phase in females, as only 9 pathways meeting our criteria were significantly enriched for rhythmic transcripts in this group. Radar plots showing the percentage of total rhythmic transcripts (p<0.05) peaking in 2-hour bins across 24 hours were constructed to examine temporal patterns in rhythmic expression. A chi square test was used to compare the distribution of transcript peak times between groups. Sequencing data will be uploaded and shared to the Gene Expression Omnibus.

### PV cell electrophysiology

For recordings under a 12:12 L:D cycle, adult male and female G42 mice were sacrificed at ZT 5-6 (light phase) or ZT 17-18 (dark phase), while for ECD experiments, female mice were sacrificed at ZT5-6 and ZT15-16 (5-6 hours after the start of either the light or dark phase in a 10:10 L:D cycle). All mice were deeply anesthetized with isoflurane and rapidly decapitated; brains were then placed into ice-cold oxygenated cutting solution consisting of (in mM): 135 *N-* methyl-D-glucamine (NMDG), 1 KCl, 1.2 KH_2_PO_4_, 1.5 MgCl_2_, 0.5 CaCl_2_, 20 choline bicarbonate, and 10 D-glucose (pH 7.4, adjusted with HCl). Sections (250µm) were cut on a vibratome (Leica VT1200S; Leica Biosystems) and placed in oxygenated modified aCSF containing (in mM): 86 NaCl, 2.5 KCl, 1.2 NaH_2_PO_4_, 35 NaHCO_3_, 20 HEPES, 25 D-glucose, 2 thiourea, 5 sodium ascorbate, 3 sodium pyruvate, 1 MgSO_4_, and 2 CaCl_2_ (pH 7.3). Sections were held at 37°C for 5 minutes before being allowed to return to room temperature for recordings.

For recordings, sections were perfused with oxygenated aCSF (∼29-32°C) containing (in mM): 119 NaCl, 2.5 KCl, 1 NaH_2_PO_4_, 26.2 NaHCO_3_, 1.3 MgCl_2_, 2.5 CaCl_2_, and 11 D-glucose; 100µM picrotoxin (Tocris Bioscience, Bristol, UK) was added to the bath to block GABA_A_ mediated transmission. The medial PFC was identified under 5x and layer 3-5 cells expressing GFP within these regions were visualized and selected for recordings using a 488nm fluorescence filter at 40x with differential interference contrast (Leica). Previous studies characterizing G42 mice have shown that all labeled PV neurons in these layers are basket cells (Miyamae et al., 2017). Borosilicate glass electrodes (∼3-5MΩ) filled with a potassium based internal solution containing (in mM): 119 KMeSO_3_, 5 KCl, 0.5 K-EGTA, 0.16 CaCl_2,_ 10 HEPES, 2.5 Mg-ATP, 0.25 Na-GTP, 12 sodium phosphocreatine, 1 L-glutathione, were used for whole cell patch clamp. Cells were held at −70mV for a 5-minute baseline prior to recording spontaneous excitatory postsynaptic currents (sEPSCs) over an additional 5 minutes. Following voltage clamp recordings, cells were switched to current clamp and the holding current was adjusted so that cells remained at approximately −70mV. A series of hyper and depolarizing current injections (500ms square pulse, −60pA to 300pA, 20pA steps) were given through the recording electrode in order to measure input resistance and evoke action potential firing. All recordings were made using a MultiClamp 700A amplifier (Molecular Devices; San Jose CA).

Voltage clamp recordings were filtered at 2kHz, amplified 5 times, and digitized at 50 kHz with a Digidata 1332 analog-to-digital converter (Molecular Devices), while current clamp recordings were filtered at 2kHz, amplified 5 times, and digitized at 10kHz. Sections were used for recordings through the end of their respective phases.

### Experimental design and statistical analyses of electrophysiological data

All analysis of PV electrophysiology recordings was performed using pClamp 11.1 (Molecular Devices). For sEPSC measurements, the first 500 events were used to assess amplitude and frequency. Current clamp recordings were analyzed for properties including: rheobase (the lowest current at which an action potential was fired) and action potential frequency. Input resistance was calculated from Ohm’s Law at a −20pA current injection.

Properties of action potentials were measured from the first action potential at rheobase. The threshold was defined as the voltage at which the rapid depolarization of the action potential began. Action potential amplitude was measured as the voltage difference between threshold and action potential peak while afterhyperpolarization amplitude was measured as the voltage difference between threshold and peak afterhyperpolarization. Half-width was measured as the action potential width at half maximal amplitude. Spike frequency adaptation was measured at a 300pA current step in cells that fired at least 4 action potentials; it was calculated as the last interspike interval divided by the first interspike interval. Responses were averaged across up to 3 repeats of the current step protocol. Exclusion criteria for recordings included poor access and cells that had a >5mV change in baseline potential in current clamp recordings.

Statistical analysis for electrophysiology recordings was performed using GraphPad Prism 9.4 (GraphPad Software, San Diego, CA, USA). As there were no sex differences in action potential frequency, phase differences in action potential frequency by current injection was assessed using a repeated measures 2-way ANOVA with sexes combined (Phase x Current injection). All other electrophysiological data under a 12:12 L:D was analyzed using a 2-way ANOVA (Sex x Phase) followed by a Sidak multiple comparisons test to reveal within sex effects of phase. In the absence of a sex effect or interaction, statistics are presented as a main effect of phase. Electrophysiological data from ECD mice were analyzed using an unpaired T-test. Data are presented as mean±SEM.

## Results

### Isolation of transcripts enriched from PV and pyramidal cells

We measured diurnal rhythms in ribosome associated RNAs, as rhythms in these RNAs are likely to reflect protein rhythms. Upon sequencing, we used principal component analysis (PCA) to confirm that the transcripts isolated from samples enriched in PV and pyramidal cells formed two separate and distinct groups based on their transcriptional profile and segregated by cell type (Fig. 1A). To further confirm the enrichment of transcripts from pyramidal cells and PV cells, we used DE analysis to measure the enrichment of cell-type markers (Fig. 1B&C).

**Figure 1.**
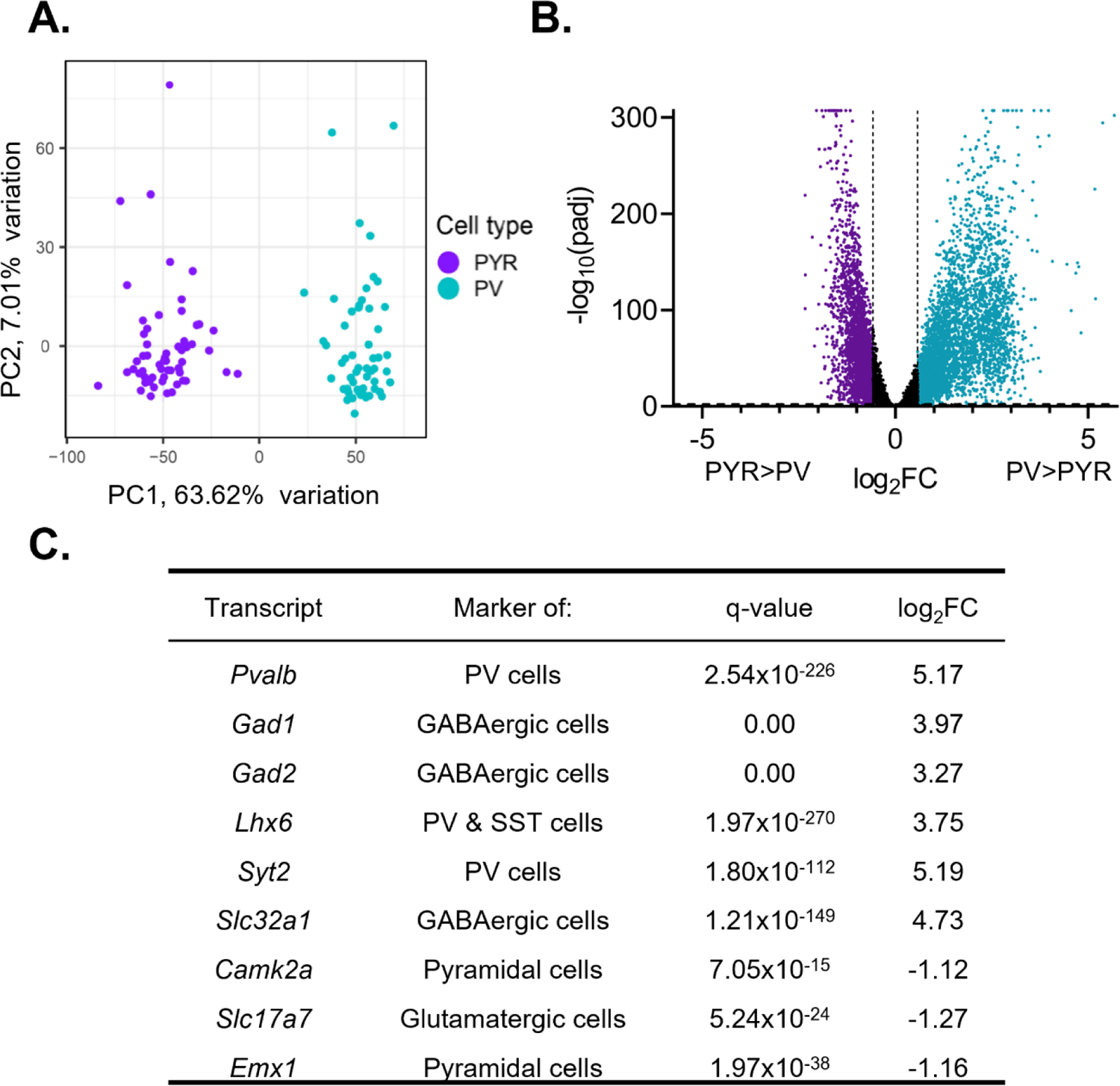
Isolated transcripts differ by cell type (A) Principal component analysis was performed using PCAtools on transcripts isolated from presumptive pyramidal cells and PV cells. Samples fall into two separate and distinct groups, segregated by cell type. (B) Differential expression analysis was performed to compare expression between PV and pyramidal cells, with transcripts from both sexes and all time points combined. Transcripts significantly enriched in pyramidal cells are colored purple while transcripts significantly enriched in PV cells are colored blue. Transcripts are considered differentially expressed (enriched) with a log_2_fold change (log_2_FC) >0.58 and FDR corrected q-value (padj) <0.01 (-log_10_(padj)>2). The log_2_FC and q-value cutoff are displayed as dashed lines. A ceiling effect is present for -log_10_(padj) in order to accommodate q-values lower than 1.0e-307. (C) The expression of known markers in PV and pyramidal cells. P-values and FDR-corrected q-values are listed. Transcripts enriched in pyramidal cells are indicated by a negative log_2_FC while transcripts enriched in PV cells are indicated by a positive log_2_FC. Canonical PV and GABAergic cell markers are enriched in PV cell samples while markers of excitatory and pyramidal cells are enriched in pyramidal cell samples. PYR=pyramidal cells, PV=parvalbumin cells, FC=fold change, padj=FDR-corrected q-value

Expression of *Pvalb* (q=2.54e-226, fold change (FC)=36.06) as well as GABAergic cell-type markers *Gad1* (q=0.00, FC=15.70), *Gad2* (q=0.00, FC=9.63), and *Slc32a1* (q=1.21e-149, FC=26.54) are significantly enriched in PV cell samples. Additionally, we find that *Syt2* (q=1.80e-112, FC= 36.53), which has been shown to be a marker of PV cells (Sommeijer and Levelt, 2012), and *Lhx6* (q=1.97e-270, FC=13.45), a marker of interneurons derived from the medial ganglionic eminence, are significantly enriched in PV cell samples relative to pyramidal cell samples. Conversely, expression of pyramidal cell markers *Camk2a* (q=7.05e-15, FC=2.17), *Slc17a7* (q=5.24e-24, FC=2.41), and *Emx1* (q=1.97e-38, FC=2.23) are significantly enriched in pyramidal cell samples. Together, this confirms that we were able to isolate transcripts selectively from either PV or pyramidal cells.

### Gene expression in pyramidal cells is highly rhythmic relative to PV cells

To identify transcripts with 24-hour rhythms in their expression, we utilized a parametric cosinor model, whereby the expression of transcripts across 24 hours is fitted to a sinusoidal curve (Cornelissen, 2014). As expected, the number of rhythmic transcripts in each sample type is dependent on the significance cutoff used to determine rhythmicity (Fig. 2A). Utilizing a cutoff of p<0.05, we find that 4525 transcripts, or ∼35% of transcripts, are rhythmic in pyramidal cells (Table 1-1), while in PV cells, 2324 (∼18%) of transcripts are rhythmic (Table 1-2). Moreover, when total rhythmic transcripts across both cell types are compared (p<0.01), 71% of all rhythmic transcripts are rhythmic exclusively in pyramidal cells, 16% are rhythmic only in PV cells, and only 13% of transcripts are rhythmic in both cell types (Fig. 2B). This highlights both the high degree of diurnal rhythmicity in transcripts from pyramidal cells relative to PV cells, as well as how distinct rhythmic transcripts are across cell types. Nevertheless, the expression of core clock components are rhythmic in both PV and pyramidal cells and known circadian genes, such as *Ciart* and *Dbp,* are found among the top 10 rhythmic transcripts in both cell types (Fig. 2C&D). Furthermore, when examining the phase and peak time of core clock genes between cell types, we find that transcript peak times overlap between cell types (Fig. 2E), suggesting that the molecular clock is conserved and oscillates in phase between excitatory and inhibitory neurons in the PFC.

**Figure 2.**
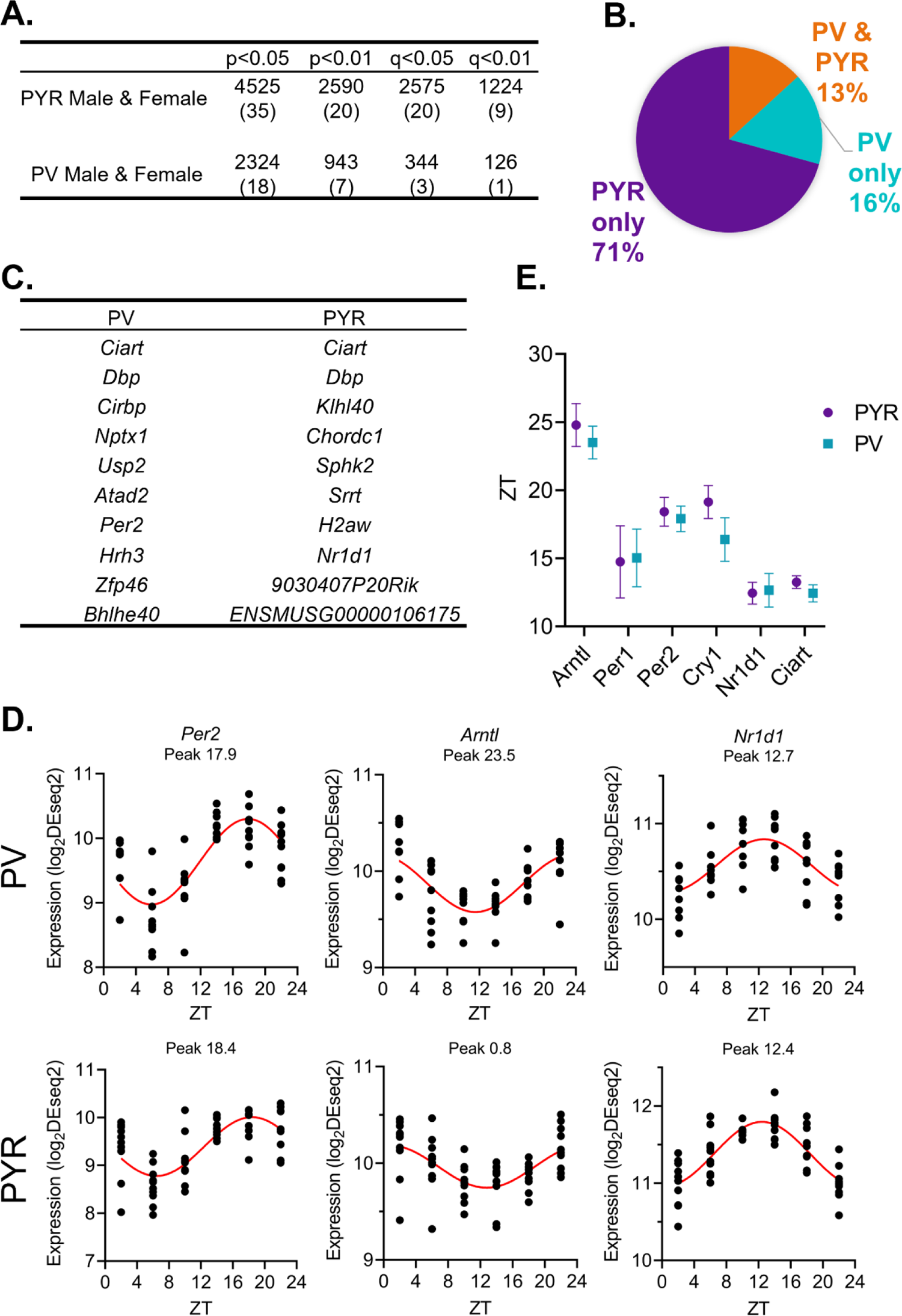
While pyramidal cells have greater transcript rhythmicity, the molecular clock is in phase across cell types (A) The number of rhythmic transcripts at different significance cutoffs in pyramidal and PV cells. The percentage of transcripts that are rhythmic are displayed in parentheses. Pyramidal cells have more rhythmic transcripts at all cutoffs. (B) The percentage of rhythmic transcripts (p<0.01) present in each cell type as a proportion of all rhythmic transcripts detected. Rhythmic transcripts are largely different between PV and pyramidal cells. (C) The top 10 rhythmic transcripts (by p-value) in PV and pyramidal cells include multiple known circadian genes. (D) Rhythmicity patterns of core clock genes in PV cells (top) and pyramidal cells (bottom). Each point represents a subject, with time of death (ZT) on the X axis and log_2_DESeq2 normalized expression on the Y axis. The fitted sinusoidal curve is shown in red. (E) The calculated peak time (ZT) and 95% confidence intervals of conserved canonical circadian transcripts, indicating that the core clock is in phase across cell types. ZT=Zeitgeber time, PYR=pyramidal cells, PV=parvalbumin cells

### Conservation in rhythmic gene expression across sexes in pyramidal cells

Previous studies have found that pyramidal cells in the mouse PFC show sex-specific diurnal rhythms in their electrophysiological properties, including diurnal variation in resting membrane potential, action potential frequency, threshold, and half-width (Roberts and Karatsoreos, 2023). However, it was unknown whether there are similar sex differences the molecular rhythms of these cells. Dividing samples by sex, we find that 2286 (∼17%) and 2097 (∼16%) transcripts are diurnally rhythmic (p<0.05) in females and males, respectively (Fig. 3A) (Tables 1-3 and 1-4). Approximately half of rhythmic transcripts in pyramidal cells are shared between sexes (1044 transcripts) (Fig.3B), significantly different from what would be expected by chance alone (Fisher’s Exact Test: p=1.33e-320; Odds Ratio: 7.79). However, as statistical power is reduced after splitting samples by sex (n=4-5/sex/timepoint in contrast to n=9-10/timepoint when sexes are combined), we also utilized a threshold-free approach (RRHO), confirming that there is a high degree of overlap in rhythmic transcripts between sexes in pyramidal cells (Fig. 3C).

**Figure 3.**
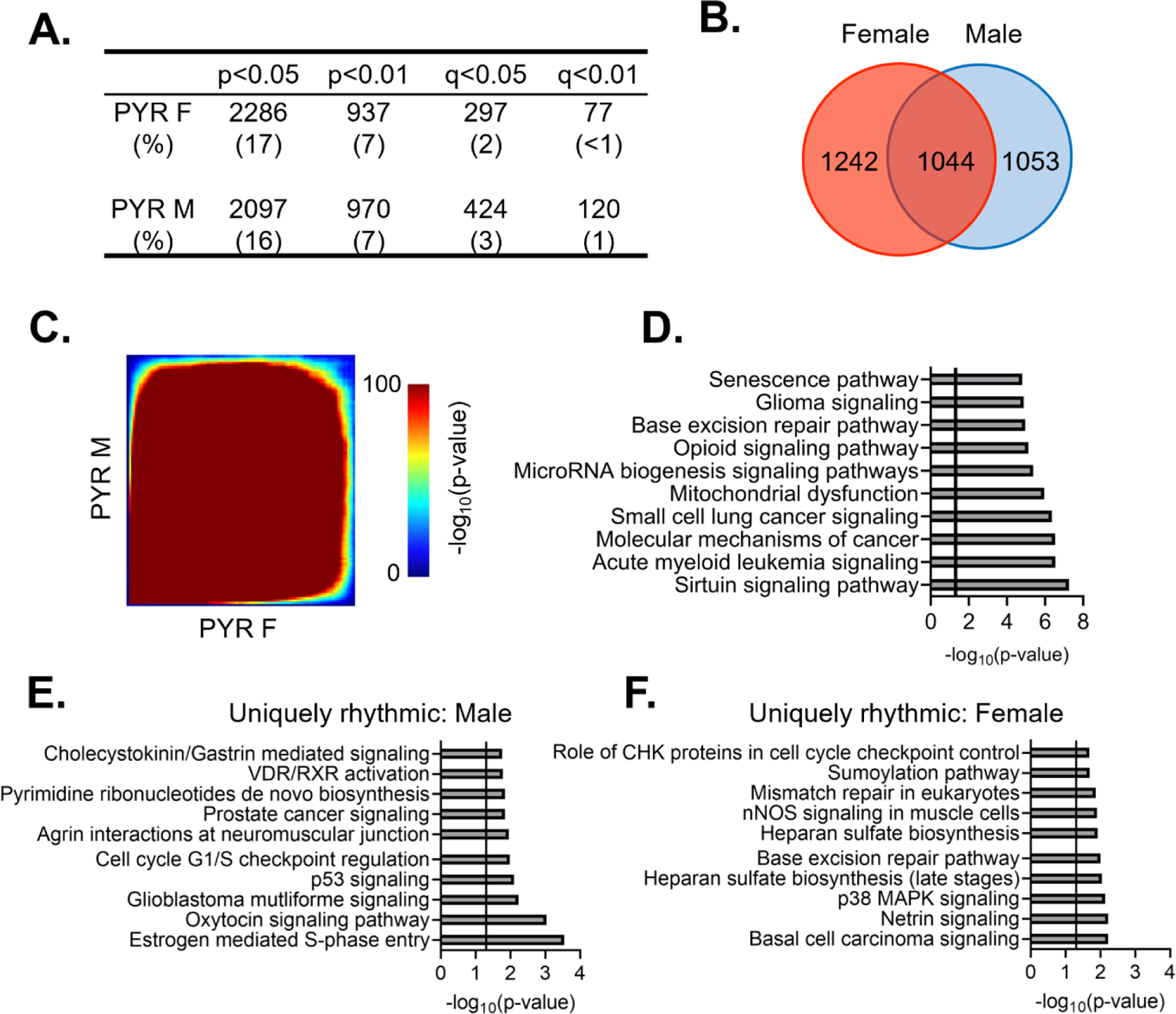
Rhythmic transcripts, and their associated processes, overlap between sexes in pyramidal cells (A) The number of rhythmic transcripts in pyramidal cells isolated from male and female animals at different significance cutoffs. The percentage of rhythmic transcripts is listed in parentheses. Both males and females show a similar number of rhythmic transcripts. (B) Venn diagram showing the overlap of rhythmic transcripts (p<0.05) between males and females. Nearly half of all rhythmic transcripts in pyramidal cells are rhythmic in both sexes. (C) Threshold-free approach (rank-rank hypergeometric overlap) assessing the overlap of rhythmicity in transcripts between males and females. There is broad overlap in rhythmic transcripts between sexes in pyramidal cells. (D) The top 10 pathways significantly enriched for rhythmic transcripts in pyramidal cells as determined by Ingenuity Pathway Analysis (Qiagen). The vertical line represents significant enrichment (-log_10_(p-value)>1.3). Top pathways include multiple cancer associated pathways. (E)&(F) The top 10 pathways enriched for transcripts that are uniquely rhythmic in either males (E) or females (F), showing sex specific roles of rhythms in pyramidal cells. PYR=pyramidal cells

The functional pathways associated with rhythmic transcripts were then assessed using Ingenuity Pathway Analysis (IPA) to determine what biological processes are most enriched for rhythmic transcripts in pyramidal cells. Given that there is extensive overlap in rhythmic transcripts between males and females, we performed this analysis with sexes combined. We find that there are multiple pathways associated with cancer among the top pathways, which include transcripts associated with apoptosis and cell division (Fig. 3D). Other pathways enriched for rhythmic transcripts are associated with oxidative phosphorylation/oxidative stress, RNA processing (microRNA biogenesis), and opioid signaling. Notably, among these pathways are transcripts coding for the NMDA receptor subunits 2A and 2B. This data suggests that rhythmic transcripts in pyramidal cells are involved in many basic cellular processes, with an important role of rhythmicity in cellular growth, energy, and survival associated processes.

While our analysis has so far revealed broad overlap of rhythmic transcripts between sexes, we wanted to determine if there are any potentially sex-specific roles of rhythmic gene expression in pyramidal cells. To ensure that we were only examining transcripts that are uniquely rhythmic in one sex, we used the following criterion: a) rhythmic at a cutoff of p<0.05 in one sex, b) not rhythmic at a cutoff of p<0.05 in the opposite sex and c) have a significant (p<0.05) ΔR^2^, indicating a significantly better goodness of fit in the same sex in which the transcript displayed rhythmicity. 337 and 271 transcripts meet this criterion in males and females, respectively (Tables 1-5 and 1-6). Similar to the combined data, uniquely rhythmic transcripts in males largely belong to pathways associated with cancer and the cell cycle (Fig. 3E). Uniquely rhythmic transcripts in males also include transcripts associated with cell adhesion (*β-catenin 1* and *laminin 1*) as well as *neuregulin 1* (agrin interactions at the neuromuscular junction). Conversely, transcripts that are uniquely rhythmic in females are primarily associated with DNA synthesis (base excision repair and mismatch repair pathways) (Fig. 3F). Pathways associated with cancer are also among the top pathways enriched for uniquely rhythmic transcripts in females, further highlighting an important role of rhythmicity in cancer associated transcripts that is conserved across sexes in pyramidal cells.

### Rhythmic transcripts in PV cells are highly sexually dimorphic

We next wanted to determine whether there are sex differences in the diurnal rhythmicity of transcripts isolated from PV interneurons. At a cutoff of p<0.05, we find that ∼14% (1819) of expressed transcripts are rhythmic in females (Table 1-7), while only ∼9% (1137) are rhythmic in males (Table 1-8)(Fig. 4A). The majority of rhythmic transcripts in PV cells, 1573 (∼86%) in females and 891 (∼78%) in males, are rhythmic only in one sex. Indeed, only 246 transcripts are rhythmic in both sexes, a finding that differs from what would be expected by chance alone (Fisher’s Exact Test: p=8.67e-14). This finding of low overlap between sexes is consistent with results from a threshold-free approach (RRHO) (Fig. 4C). Notably, the overlap in rhythmic transcripts between sexes in PV cells is lower than that of pyramidal cells (Odds Ratio 1.82 in PV cells versus 7.79 in pyramidal cells).

**Figure 4.**
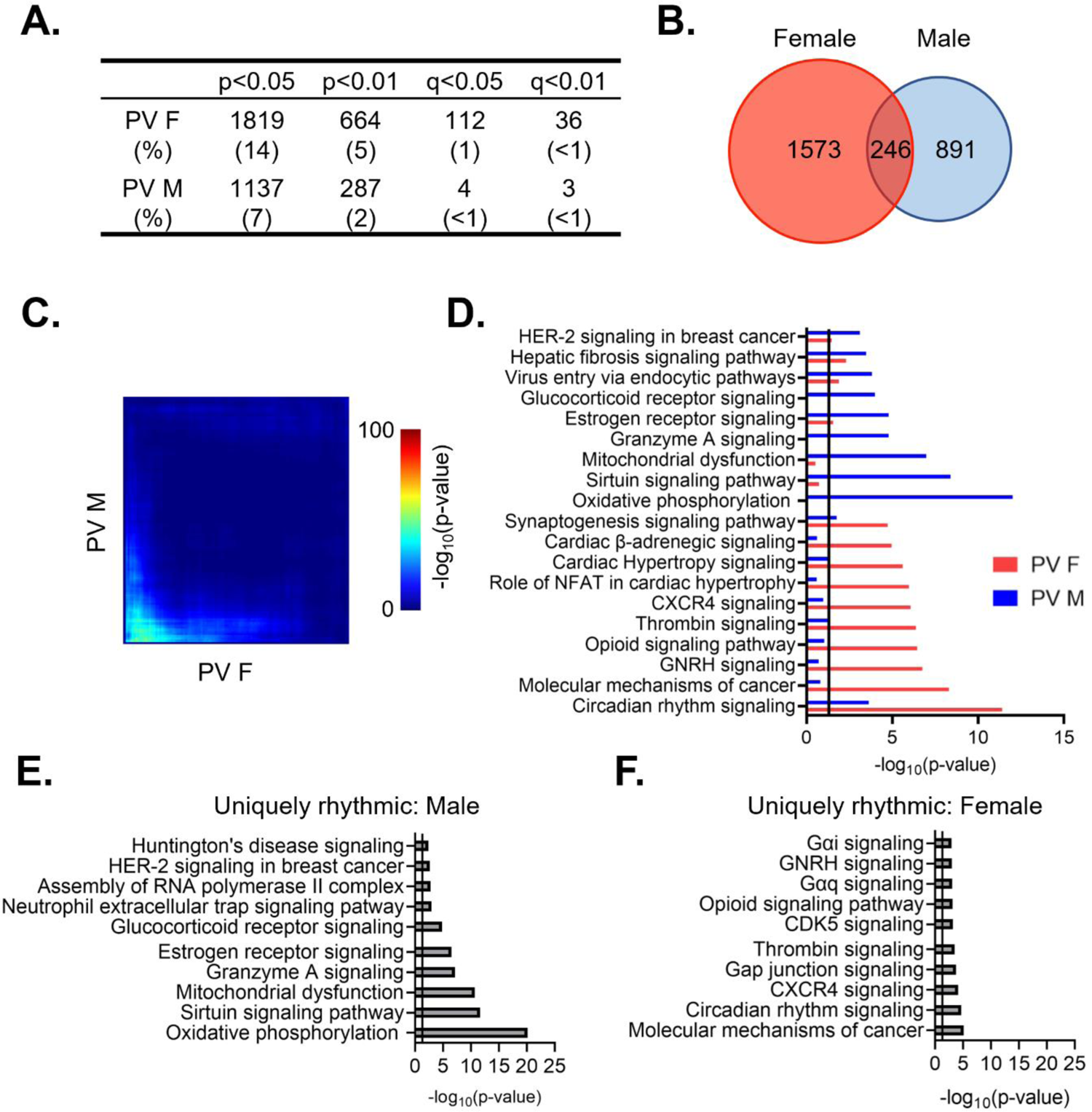
Rhythmic transcripts from PV cells are largely distinct between sexes (A) The number of rhythmic transcripts in PV cells isolated from male and female animals at different cutoffs. Females have more rhythmic transcripts at all significance cutoffs. (B) Venn diagram showing the overlap of rhythmic transcripts (p<0.05) between males and females. Few transcripts are rhythmic in both males and females. (C) Threshold-free approach (rank-rank hypergeometric overlap) assessing the overlap of rhythmic transcripts (regardless of p-value) between males and females. Very little overlap confirms that rhythmic transcripts are largely distinct in males and females. (D) The top 10 enriched rhythmic pathways in males and females as determined by Ingenuity Pathway Analysis (Qiagen). Pathways associated with rhythmic transcripts are largely different between males and females. (E)&(F) The top 10 pathways enriched for transcripts that are uniquely rhythmic in either males (E) or females (F), showing sex specific roles of rhythms in PV cells. PV=parvalbumin cells

To determine whether the functions associated with rhythmic transcripts in PV cells also differ between males and females, we compared the top enriched pathways as determined by IPA. We find that circadian rhythm signaling as well as estrogen receptor signaling and synaptogenesis signaling are significantly enriched for rhythmic transcripts in both males and females, although the transcripts within each pathway largely differ by sex (Fig. 4D). Similar to our findings in pyramidal cells, pathways involving cancer are among the top enriched pathways in both sexes, suggesting an important role of rhythmicity in proliferation and the cell cycle that is consistent across cell types. However, many enriched pathways in PV cells are exclusive to one sex. In males, rhythmic transcripts are predominantly associated with mitochondria and metabolism, as well as vesicular trafficking (virus entry via endocytic pathways). Conversely, in females, many of the top pathways represented by rhythmic transcripts involve intercellular communication and synaptic transmission (opioid signaling, β-adrenergic signaling, and hormone signaling), with rhythmic transcripts within these pathways largely associated with G-protein coupled receptor (GPCR) mediated signaling. This suggests that there are sex-specific functions of rhythmic transcripts in PV cells.

We further investigated these sex differences in PV cells by determining the functional pathways associated with transcripts that are uniquely rhythmic in one sex. Using the same criteria as described above, transcripts that are uniquely rhythmic in males (467 transcripts) (Table 1-9) belong to pathways associated with transcription as well as oxidative phosphorylation and metabolism (Fig. 4E). Notably, oxidative phosphorylation is very highly enriched for transcripts that are uniquely rhythmic in males, including many transcripts coding for subunits of the electron transport chain. Transcripts that are uniquely rhythmic in females (929 transcripts) (Table 1-10) are associated with GPCR mediated signaling, including many transcripts belonging to the adenylate cyclase gene family (Fig. 4F). This demonstrates that transcripts that are uniquely rhythmic in only one sex are involved in very different biological processes and highlights the unique role of rhythmic gene expression in mitochondrial associated processes in PV cells from males.

### Temporal patterns of rhythmic transcript expression in pyramidal cells differ between sexes

To assess broad temporal patterns in rhythmic gene expression, we plotted the percentage of rhythmic transcripts (p<0.05) by their peak time (ZT) across 24 hours by sex. In pyramidal cells, we find that the distribution of transcript peak times differs between sexes (X^2^ (df=10; n=2097-2286)=257.87, p=1.20e-49). In females, rhythmic transcripts primarily peak at distinct time points, forming two groups of transcripts with similar peak times (Fig. 5A). One group of transcripts peaks in the light phase at ZT4-7 and consists of ∼29% of rhythmic transcripts, while another group, made up of ∼26% of rhythmic transcripts, peaks in the dark phase at ZT16-19. The remaining transcripts peak in small groups across 24 hours (<10% per two-hour bin), with the fewest transcripts peaking around the transition between the light and dark phases (ZT12-13). In contrast, the peak times in males are fairly evenly distributed across 24 hours, with each two-hour bin representing ∼6-10% of rhythmic transcripts.

**Figure 5.**
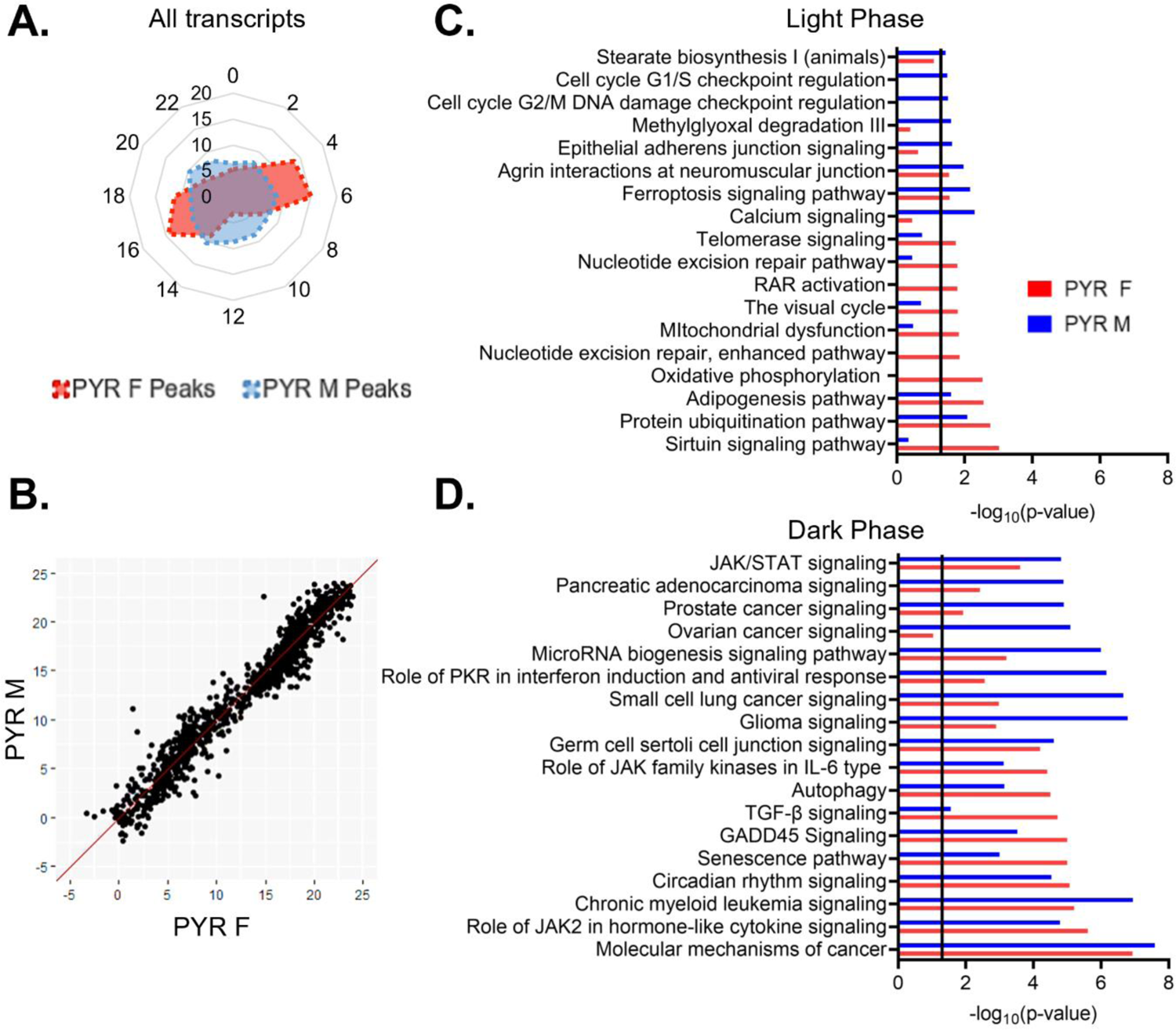
Temporal patterns of rhythmic transcript expression in pyramidal cells differ between sexes (A) Radar plot showing the peak time of rhythmic transcripts (p<0.05) across a 24-hour period (ZT). Data is presented as the proportion of total rhythmic transcripts peaking within each 2-hour bin. Females show two distinct groups of rhythmic transcripts, one peaking in the light phase and the other peaking in the dark. Conversely, peak times in males are relatively evenly distributed across 24 hours. (B) Phase concordance plot showing the peak time (ZT) of individual shared rhythmic transcripts in females (X axis) and males (Y axis). Shared rhythmic transcripts peak at similar times in both males and females. (C)&(D) The top 10 pathways (IPA) enriched for rhythmic transcripts in pyramidal cells by the phase in which they peak. Transcripts that peak during the light phase (ZT0-12) are plotted in (C) while transcripts that peak during the dark phase (ZT12-24) are plotted in (D). There is more overlap between sexes, and greater enrichment for rhythmic transcripts, in the dark phase. ZT=Zeitgeber time, PYR=pyramidal cells, IPA=Ingenuity Pathway Analysis

To determine whether these sex differences are driven by transcripts that are only rhythmic in one sex or due to the same transcripts peaking at different times of day in males and females, we next examined the phase concordance of all rhythmic transcripts that are shared between males and females, with the peak time in females on the X axis and the peak time in males on the Y axis (Fig. 5B). The majority (∼92%) of transcripts peak within 3 hours in males and females, indicating that shared rhythmic transcripts generally peak in phase between sexes.

Rhythmic transcripts that peak during the light or the dark phase may be involved in biological processes that are specifically timed to the animal’s inactive or active phase, respectively. Therefore, to determine the functional pathways represented by rhythmic transcripts in each phase, we split rhythmic transcripts in pyramidal cells by the phase in which their expression peaks (“light (inactive) phase” ZT0-12, “dark (active) phase” ZT12-24). As we observed different temporal patterns in the peak times of rhythmic transcripts between males and females, we performed this analysis on samples split by sex. During the light phase, four of the top enriched pathways show significant enrichment for rhythmic transcripts in both sexes (protein ubiquitination, adipogenesis, ferroptosis signaling, and agrin interactions at neuromuscular junction) (Fig 5C). Broadly, in males, rhythmic transcripts that peak during the light phase are associated with calcium signaling, protein processing (protein ubiquitination), the cell cycle, and the cytoskeleton (agrin interactions at the neuromuscular junction and epithelial adherens junction signaling). In females, rhythmic transcripts that peak during the light phase are also associated with protein processing (protein ubiquitination), as well as oxidative phosphorylation and DNA replication and repair (nucleotide excision repair). In the dark phase, pathways generally show a higher enrichment for rhythmic transcripts than pathways in the light phase (Fig. 5D). Indeed, many of the pathways in the dark phase match the top pathways when peak time is not considered, suggesting that transcripts that peak during the dark phase drove that analysis. All pathways that fall within the top enriched pathways in one sex are also significantly enriched in the other sex, with the exception of ovarian cancer signaling, which is one of the top pathways in males. Broadly, transcripts peaking during the dark phase in pyramidal cells are associated with cancer and cytokine signaling in both sexes. These data suggest that during the active (dark) phase, the biological processes associated with rhythmic transcripts are widely conserved across sex, while during the inactive (light) phase, there are more sex-specific roles of rhythmic transcripts in pyramidal cells.

### Temporal patterns of rhythmic gene expression and associated processes differ in PV cells

When we examine the temporal relationship of rhythmic transcripts in PV cells, we find that the distribution of transcript peak times differ by sex (X_2_ (df=10, n=1137-1819)=208.45, p=2.77e-39) (Fig. 6A). In females, transcripts primarily peak at two time points, with one group peaking in the light phase (ZT6-9), made up of ∼38% of rhythmic transcripts, and one group peaking in the dark phase (ZT18-21), made up of ∼30% of rhythmic transcripts. In males, there is a group of rhythmic transcripts that peak during the light phase at ZT6-11 (∼35% of transcripts), partially overlapping with the group of rhythmic transcripts that peak during the light phase in females. However, in the dark phase, transcript peak times in males are fairly evenly distributed, with <10% of transcripts peaking in each 2-hour bin. However, when only transcripts that are rhythmic in both sexes are examined, most transcripts peak around the same time, with 80% peaking within 3 hours across sexes (Fig. 6B).

**Figure 6.**
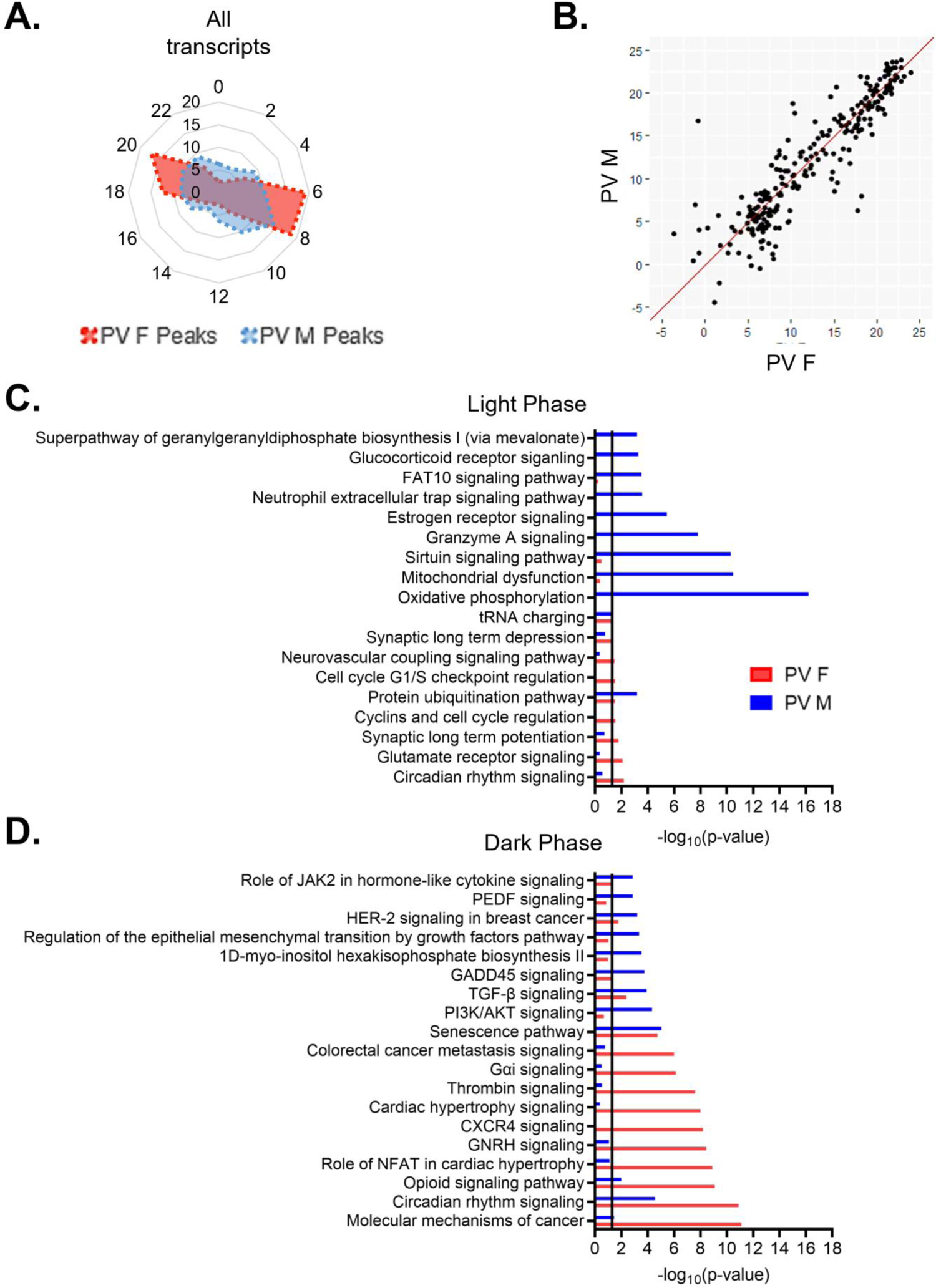
PV cells show more overlap in biological processes enriched for rhythmic transcripts between sexes in dark phase (A) Radar plot showing the peak time of rhythmic transcripts (p<0.05) in PV cells from both males and females. Data is presented as the proportion of total rhythmic transcripts peaking within 2-hour bins across a 24-hour period (ZT). Rhythmic transcripts in females form two groups, one peaking in the light phase and one peaking in the dark phase. In males, rhythmic transcripts form a small group that peaks in the light phase, but are fairly evenly distributed in the dark phase. (B) Phase concordance plot showing the peak time (ZT) of individual shared rhythmic transcripts in females (X axis) and males (Y axis). Shared rhythmic transcripts largely peak at similar times between sexes. (C)&(D) The top 10 pathways (IPA) enriched for rhythmic transcripts in males and females separated by the phase in which they peak. Transcripts that peak during the light phase (ZT0-12) are plotted in (C) while transcripts that peak during the dark phase (ZT12-24) are plotted in (D). There is more overlap in pathways enriched for rhythmic transcripts in the dark phase. ZT=Zeitgeber time, PV=parvalbumin cells, IPA=Ingenuity Pathway Analysis

We next investigated whether the biological processes enriched for rhythmic transcripts show phase specific overlap between sexes in PV cells. We find that although there is partial overlap in the temporal patterns of rhythmic transcripts during the light phase, the top functional pathways associated with these transcripts are vastly different (Fig. 6C). The only top pathway that is significantly enriched for rhythmic transcripts in both sexes during the light phase is protein ubiquitination, suggesting a conserved role of rhythmic transcripts in protein processing. Many of the pathways associated with transcripts that peak during the light phase in males match the top pathways identified when transcript peak times are not considered, including oxidative phosphorylation and mitochondrial dysfunction. In females, pathways enriched for rhythmic transcripts that peak during the light phase are largely associated with intercellular communication, including glutamate receptor signaling and long-term synaptic potentiation/depression. Notably, there are only 9 pathways significantly enriched for rhythmic transcripts in the light phase in females. In the dark phase, 7 of the top pathways are enriched for rhythmic transcripts in both sexes, including those associated with circadian rhythm signaling and cancer. In males, rhythmic transcripts are enriched for processes associated with cell survival/aging (senescence) and cytokine signaling, while rhythmic transcripts in females are associated with GPCR mediated signal transduction (Fig. 6D). Together, these data suggest that while the molecular clock may be synchronized between PV and pyramidal cells, there are broad cell-type and sex-specific differences in transcript rhythmicity and their associated biological processes, as summarized in Fig. 7.

**Figure 7.**
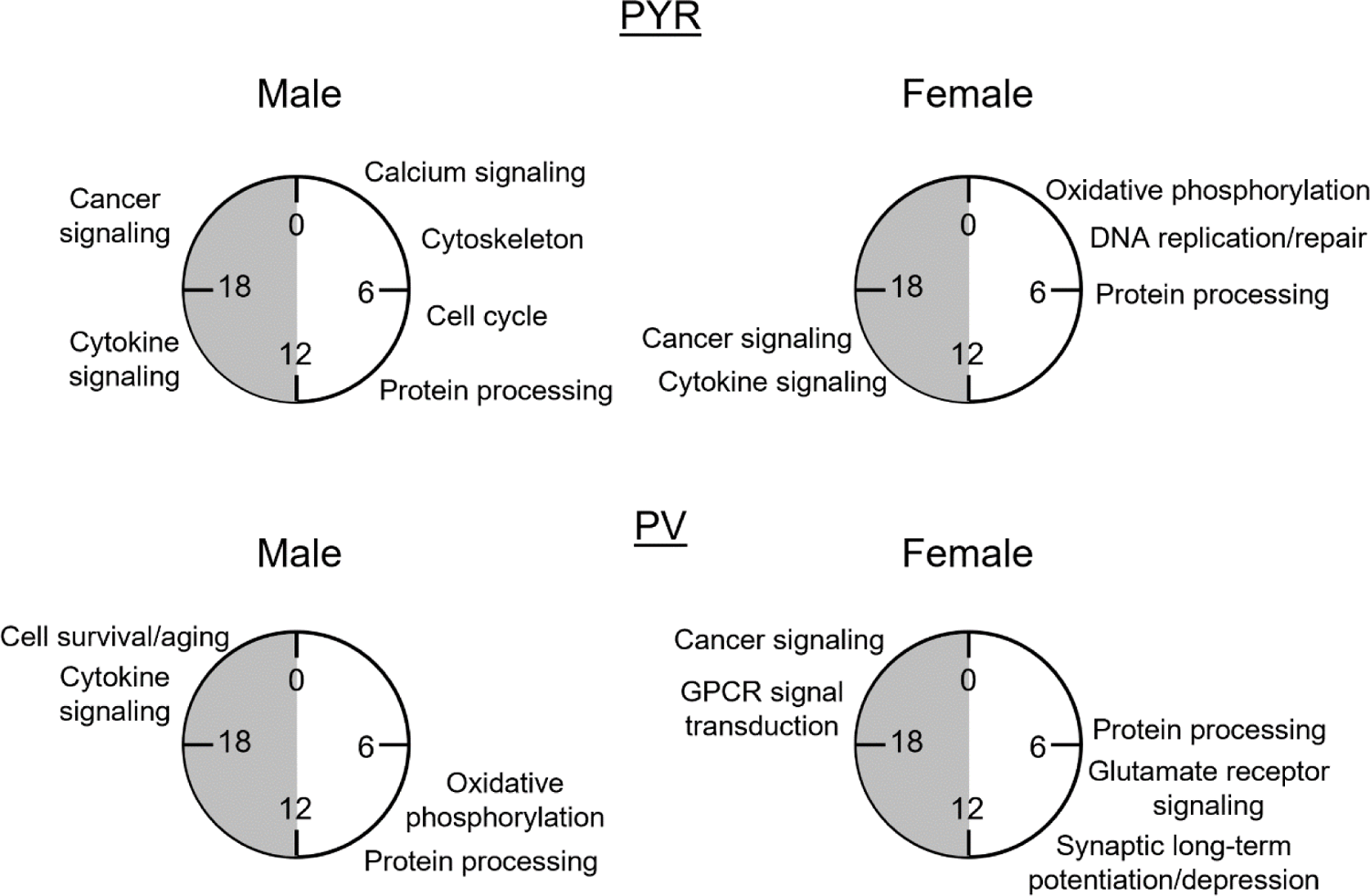
The biological processes associated with rhythmic transcripts differ by sex and cell type. Summary of key processes associated with rhythmic (p<0.05) transcripts in pyramidal cells (top) and PV cells (bottom) by the phase in which they peak. Processes are approximately aligned with the peak time (ZT) of transcripts in each phase as shown in radar plots in figures 5 and 6. This figure highlights the unique role of rhythms in each sex and cell type. PYR=pyramidal cells, PV=PV cells, GPCR=G-protein coupled receptor, ZT=Zeitgeber time

### Molecular rhythms are more similar across cell types in females than males

To assess overlap across cell types, we next used RRHO to compare rhythmic transcripts between PV and pyramidal cells. In females, we find that there is moderate overlap in rhythmic transcripts (Fig. 8A). Of note, the observed overlap across cell types in females is higher than the overlap observed within PV cells between sexes, as shown previously in Fig. 4C. Moreover, the functional pathways associated with rhythmic transcripts were assessed to look for pathways conserved across cell types and those unique to each individual cell type. In females, there is broad overlap among the top enriched pathways in PV and pyramidal cells; these overlapping pathways include autophagy, molecular mechanisms of cancer, and circadian rhythm signaling (Fig. 8B). To investigate possible cell-type specific differences in rhythmicity, we examined transcripts that are uniquely rhythmic in one cell type, as defined previously. In females, uniquely rhythmic transcripts in pyramidal cells (1244 transcripts) (Table 1-11) belong to pathways involved in processes such as cytokine signaling, translation (tRNA charging), axonal migration (ephrin A signaling), and DNA repair (base excision repair) (Fig. 8C (top)). In contrast, rhythmic transcripts unique to PV cells in females (848 transcripts) (Table 1-12) belong to pathways involved in intercellular communication, including the synaptogenesis signaling pathway and GABA receptor signaling, as well as transcriptional regulation (DNA methylation and transcriptional repression) (Fig. 8C (bottom)). This suggests a cell-type specific role of rhythmic gene expression in neuronal communication that is unique to PV cells in females.

**Figure 8.**
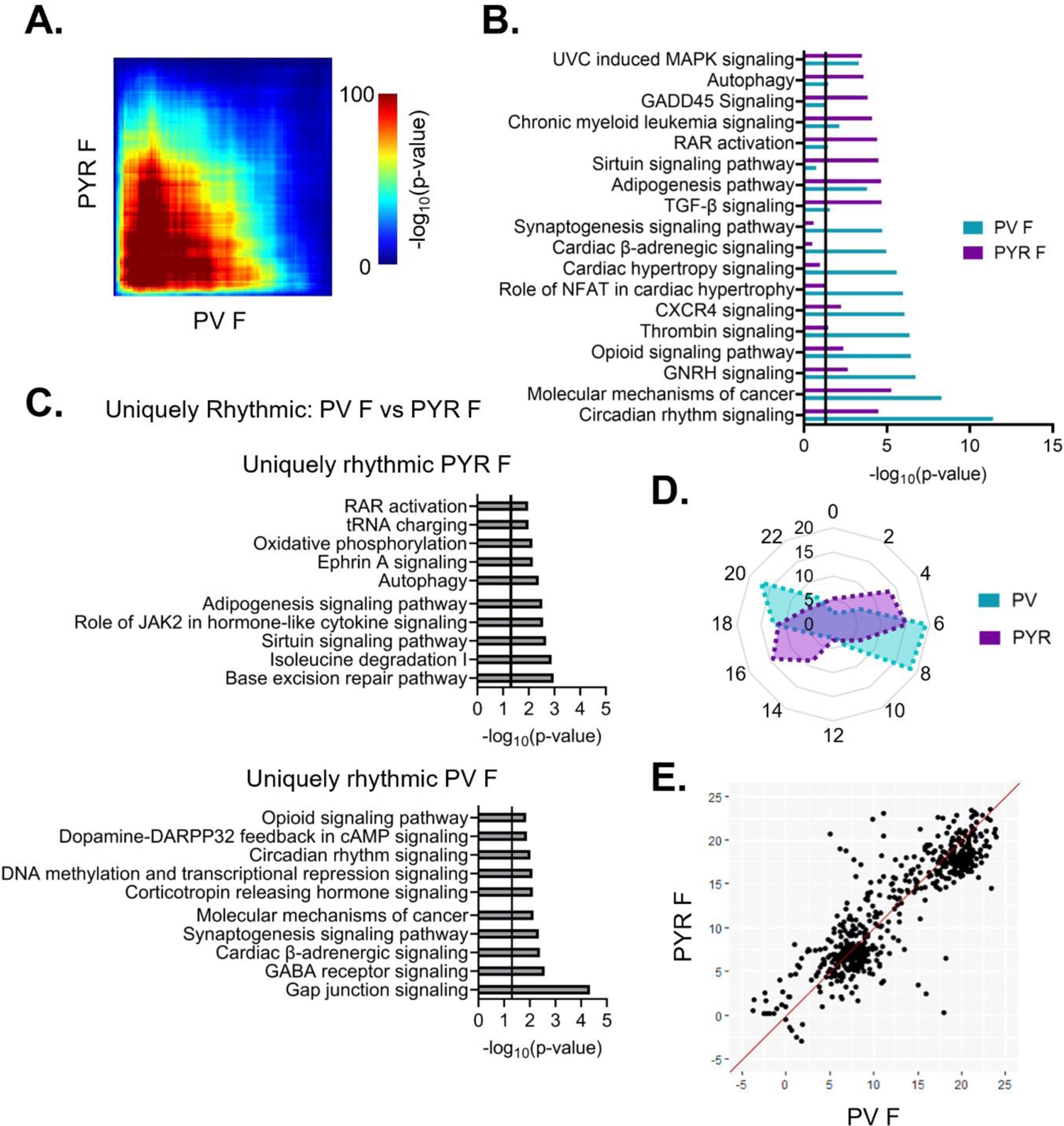
Shared patterns of transcript rhythmicity across cell types in females (A) Threshold-free approach (rank-rank hypergeometric overlap) assessing the overlap of rhythmic transcripts between cell types in females. There is moderate overlap in rhythmic transcripts between cell types in females. (B) Pathway analysis (IPA) of rhythmic transcripts (p<0.05) across cell types in females. The top 10 pathways and their overlap between cell types are plotted. Pathways enriched for rhythmic transcripts show substantial overlap between cell types particularly in pathways associated with cancer. (C) The top 10 pathways (IPA) associated with transcripts that are uniquely rhythmic in pyramidal cells (top) or PV cells (bottom). Uniquely rhythmic transcripts in pyramidal cells are associated with a variety of processes including cytokine signaling, autophagy, and DNA repair while uniquely rhythmic transcripts in PV cells are largely associated with intercellular communication. (D) Radar plot comparing the peak times (ZT) of rhythmic transcripts between cell types across 24 hours. Data is presented as the percentage of total rhythmic transcripts peaking within each 2-hour bin. In both cell types, females show two distinct groups of transcripts, one peaking in the light phase and the other during the dark phase, with a phase shift of 2-4 hours between cell types. (E) Phase concordance plots showing the peak time (ZT) of individual shared rhythmic transcripts in PV cells (X axis) and pyramidal cells (Y axis). Shared rhythmic transcripts tend to peak around the same time between cell types. IPA=Ingenuity Pathway Analysis, PV=parvalbumin cells, PYR=pyramidal cells, ZT=zeitgeber time

We next compared the temporal patterns of all rhythmic transcripts to determine if overall rhythmic gene expression is synchronized in females across cell types. Plotting the peak time of rhythmic transcripts across 24 hours reveals that, in females, the distribution of transcript peak times differs between cell types (Χ^2^ (df=10, n=1819-2286)=512.88, p=7.79e-104) (Fig 8D). Nevertheless, both pyramidal cells and PV cells show two distinct groups of transcripts one that peaks during the light phase and one that peaks during the dark phase. However, the timing of these groups differs by approximately 2-4 hours, with rhythmic transcripts peaking earlier in pyramidal cells than in PV cells. When examining the phase concordance of individual transcripts that are rhythmic in both PV and pyramidal cells from females, peak timing is relatively consistent between cell types, with ∼70% of transcripts peaking within 3 hours between cell types (Fig. 8E). Unlike in the radar plot, there is not a clear shift in females whereby peak timing is earlier in pyramidal cells than PV cells.

We next performed the same analysis to compare rhythms across cell types in males. Unlike in females, we find minimal overlap in rhythmic transcripts between cell types in males using RRHO (Fig. 9A). Additionally, while most pathways enriched for rhythmic transcripts do not overlap, those that do are primarily associated with circadian rhythm signaling and cancer (Fig. 9B). When we investigate cell-type differences in males by examining transcripts that are uniquely rhythmic in either PV or pyramidal cells, we find that transcripts that are uniquely rhythmic in pyramidal cells (1390 transcripts) (Table 1-13) are largely enriched in processes such as cell division and proliferation, with many cancer and cell cycle associated pathways among the top enriched pathways (Fig. 9C (top)). Conversely, uniquely rhythmic transcripts from PV cells in males (581 transcripts) (Table 1-14) are particularly enriched in processes associated with oxidative phosphorylation and metabolism, as well as other basic biosynthetic processes (assembly of RNA polymerase and cysteine biosynthesis) (Fig. 9C (bottom)).

**Figure 9.**
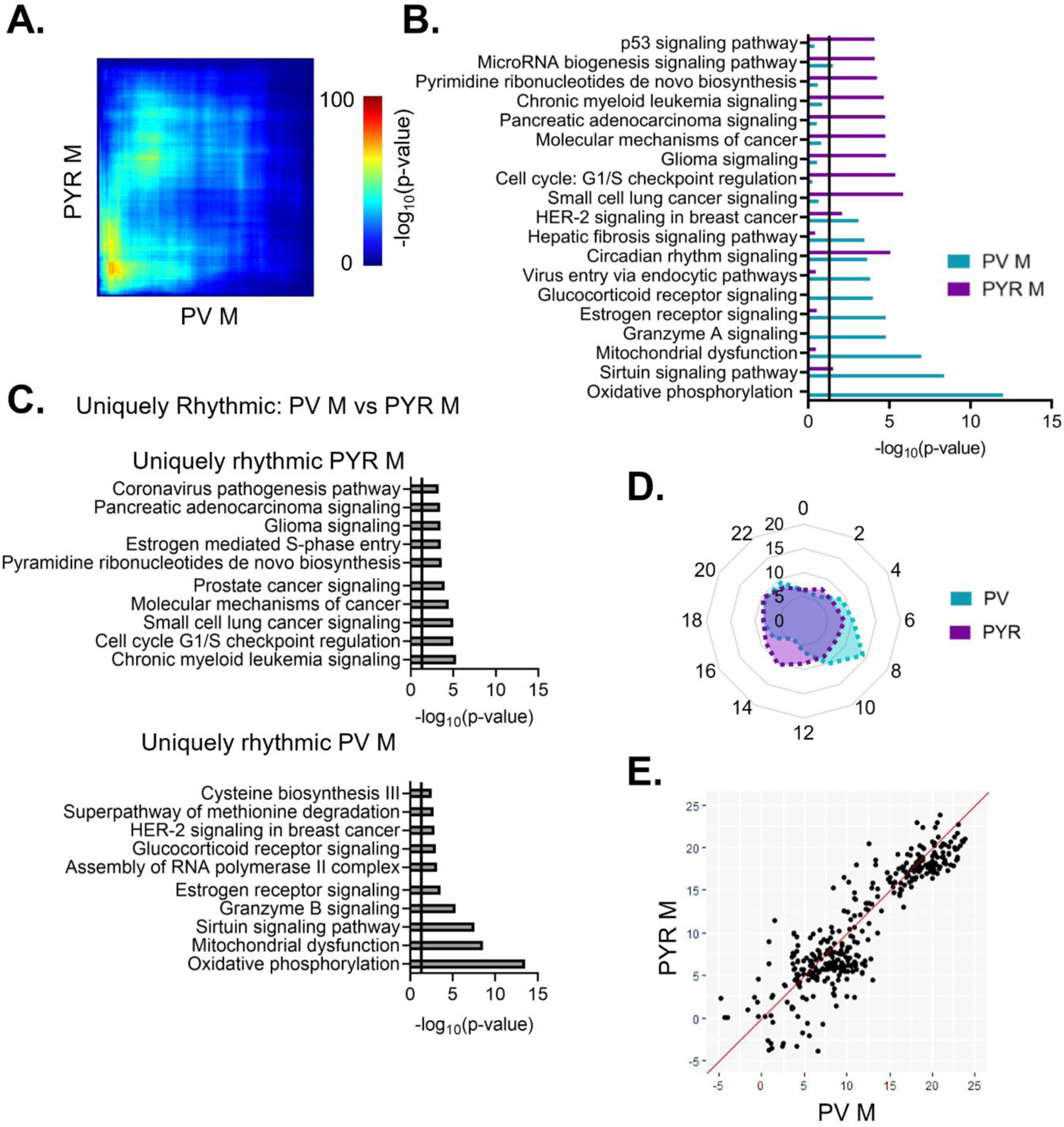
Rhythmic transcripts are largely cell-type specific in males (A) Threshold-free approach (rank-rank hypergeometric overlap) assessing the overlap of rhythmic transcripts between PV and pyramidal cells in males. There is limited overlap in rhythmic transcripts between cell types in males. (B) Pathway analysis (IPA) of rhythmic transcripts (p<0.05) across cell types in males. The top 10 pathways and their overlap are plotted. There is very little overlap in pathways enriched for rhythmic transcripts between cell types in males. (C) Top pathways (IPA) enriched for transcripts that are uniquely rhythmic in pyramidal cells (top) or PV cells (bottom). Uniquely rhythmic transcripts are largely associated with the cell cycle in pyramidal cells and oxidative phosphorylation in PV cells. (D) Radar plot showing the peak time (ZT) of rhythmic transcripts across 24 hours, presented as the percentage of total rhythmic transcripts peaking within each 2-hour bin. In males, rhythmic transcripts peak relatively evenly across 24 hours, with a small cluster peaking in the light phase in PV cells. (E) Phase concordance plot showing the peak time (ZT) of individual transcripts in PV cells (X axis) and pyramidal cells (Y axis). Shared rhythmic transcripts largely peak within 3 hours of one another. IPA=Ingenuity Pathway Analysis, PV=parvalbumin cells, PYR=pyramidal cells, ZT=zeitgeber time

When examining the temporal relationship of rhythmic transcripts between PV and pyramidal cells in males, we find that the distribution of transcript peak times differs by cell type (Χ^2^ (df=10, n=1137-2097)=85.89, p=3.48e-14) (Fig. 9D). However, when the phase concordance of transcripts that are rhythmic in both cell types is examined, the majority (68%) of shared rhythmic transcripts peak within 3 hours between cell types, similar to our findings in females (Fig. 9E). Together, these findings suggest that there is not a consistent phase shift in the timing of rhythmic gene expression between PV and pyramidal cells, with shared rhythmic transcripts showing high phase concordance in both sexes.

### Diurnal changes in PV cell electrophysiology are sex-specific in the PFC

While rhythms in actively translated mRNAs provide valuable insight into how time of day may broadly affect cellular processes, electrophysiological measurements are crucial for understanding the output of these rhythms. Previous studies have found a sex-specific time of day effect on the electrophysiological properties of pyramidal cells in the PFC, including diurnal changes in resting membrane potential, firing frequency, action potential threshold, half-width, and membrane conductance (Roberts and Karatsoreos, 2023). However, it remains unknown if there are similar diurnal differences in the electrophysiological properties of PV cells in the PFC. Indeed, as we find that there are rhythms in transcripts coding for voltage gated ion channels and associated proteins in PV cells (e.g. *Kcnq2* in females and *Kcnip2* in males), we predicted that there may be rhythms in PV cell excitability. Therefore, we used G42 mice, which express GFP in PV interneurons in the cortex, to measure the excitability and action potential properties of PV cells in the mouse PFC during both the light and the dark phase. In response to a series of depolarizing current steps, we find that there is no difference in either the action potential frequency (Main effect of phase: F(1,34)=0.09, p=0.77; current x phase interaction: F(12,408)=0.03, p>0.99) (Fig. 10A&B) or in the rheobase (Main effect of phase: F(1,32)=0.81, p=0.37 (Fig. 10C) between cells recorded in the light and dark phase. Furthermore, we find no difference in input resistance between PV cells recorded in the light or dark phase (Main effect of phase: F(1,32)=0.47, p=0.50) (Fig. 10D). Together, this data indicates that phase does not have an impact on the intrinsic excitability of PV cells in the PFC.

**Figure 10.**
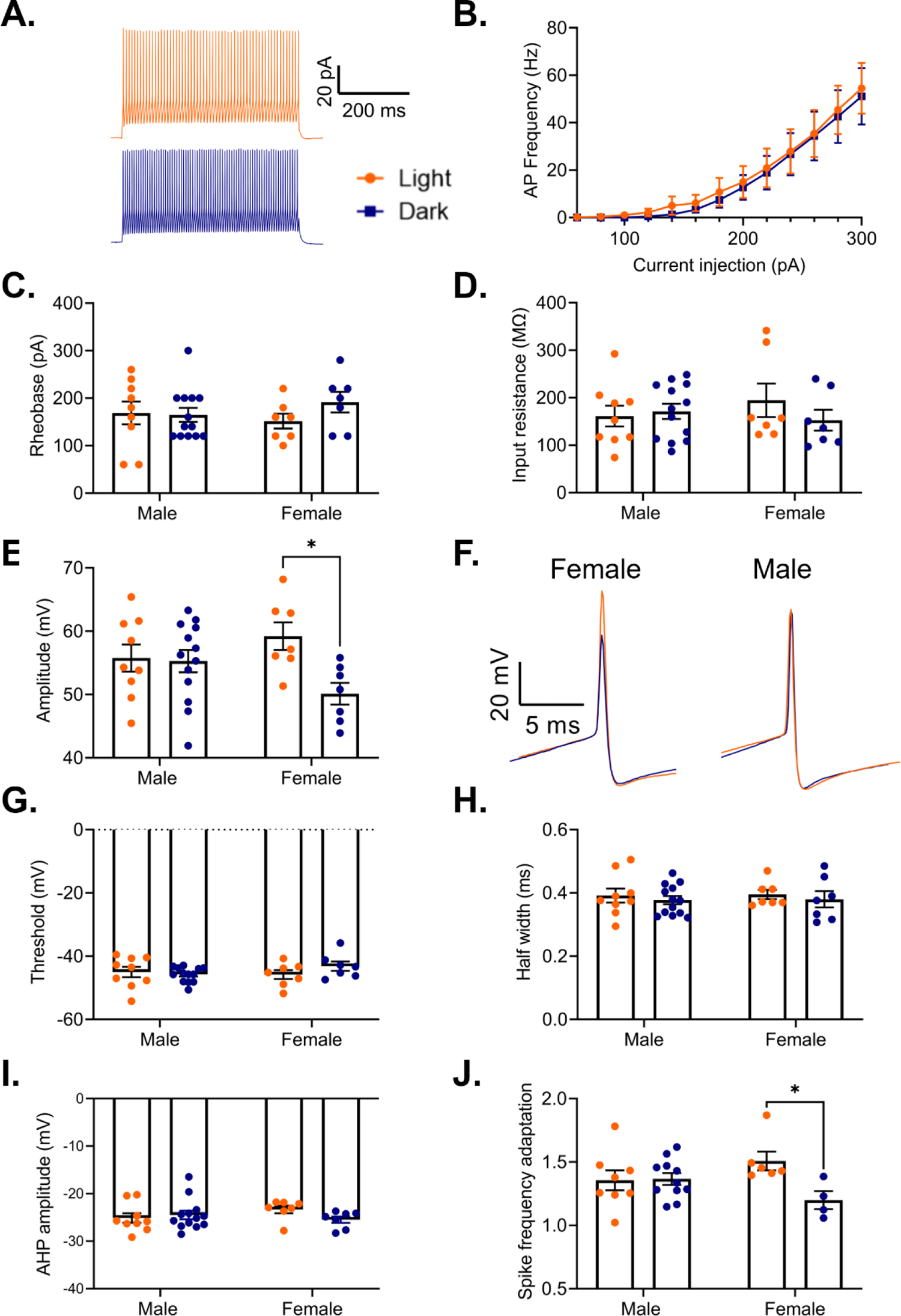
Sex-specific effects of phase on PV interneuron electrophysiology (A) Representative traces showing action potential firing of a PV cell in response to a 500ms 300pA current pulse in the light phase (orange) and the dark phase (blue). (B) Frequency of action potentials evoked from PV cells in response to current steps during the light phase and the dark phase. There is no effect of phase on action potential frequency (Main effect of phase: F(1,34)=0.00, p=0.77; Current x Phase interaction: F(12,408)=0.03, p>0.99). (C) There is no difference in the rheobase of PV cells recorded in the light and dark phases (Main effect of phase: F(1,32)=0.81, p=0.37). (D) There is no difference in the input resistance of PV cells recorded in the light and dark phases (Main effect of phase: F(1,32)=0.47, p=0.50). (E) There is a significant sex by phase interaction on action potential amplitude (Sex x Phase interaction: F(1,32)=4.41, p=0.04). In females, action potentials have a significantly higher amplitude in the light phase compared to the dark phase (p=0.01). (F) Representative traces showing a diurnal difference in action potential amplitude in females (left). No difference is observed in males (right). (G) There is no effect of phase on action potential threshold (Main effect of phase: F(1,32)=0.56, p=0.46). (H) There is no effect of phase on action potential half-width (Main effect of phase: F(1,32)=0.60, p=0.44). (I) There is no effect of phase on afterhyperpolarization amplitude (Main effect of phase: F(1,32)=0.65, p=0.43). (J) There is a significant sex by phase interaction on spike frequency adaptation (last interspike interval/first interspike interval) (Sex x Phase interaction: F(1,25)=5.00, p=0.03). Females show less spike frequency adaptation during the dark phase (p=0.03). Panel (B) n=16-20 cells/group, 8-9 animals/group, panels (C-I) n=7-13 cells/group, 3-5 animals/group, panel (J) n=4-11 cells/group, 2-5 animals/group. *p<0.05, PV=parvalbumin cells, AHP=afterhyperpolarization

We next assessed the evoked action potentials for diurnal changes in action potential features. Here, we find a significant sex by phase interaction on action potential amplitude (Sex x Phase interaction: F(1,32)=4.41, p=0.04), with post-hoc testing revealing that in females, action potentials during the dark phase have a significantly lower peak amplitude than action potentials during the light phase (p=0.01) (Fig. 10E&F). However, we find no effect of phase on action potential threshold (Main effect of phase: F(1,32)=0.56; p=0.46) (Fig. 10G), action potential half-width (Main effect of phase: F(1,32)=0.60, p=0.44)(Fig.10H), or afterhyperpolarization amplitude (Main effect of phase: F(1,32)=0.65; p=0.43) (Fig. 10I). We next determined whether there is a diurnal change in the characteristic ability of PV cells to fire rapid, non-adapting trains of action potentials by measuring spike frequency adaptation, defined as the interspike interval between the last two action potentials in a train divided by the interspike interval between the first two action potentials. Analysis reveals a significant interaction (Sex x Phase interaction: F(1,25)=5.00, p=0.03), suggesting that both sex and time of day play a role in this canonical feature of PV cells (Fig. 10J). Post-hoc testing reveals that in females, PV cells in the dark phase display a lower spike frequency adaptation than those in the light phase (p=0.03), suggesting a sex-specific enhanced ability to sustain high frequency firing in the dark phase.

As previous studies found that pyramidal cells in males show increased excitability during the light phase, with increased action potential frequency and a decreased threshold (Roberts and Karatsoreos, 2023), we next examined whether those diurnal changes affect the total excitatory drive to PV cells (Fig. 11A). However, we find no significant effect of phase on either frequency (Main effect of phase: F(1,24)=0.003, p=0.96) or amplitude (Main effect of phase: F(1,24)=0.10, p=0.76) of sEPSCs onto PV cells in males or females (Fig. 11B). This suggests that while there are diurnal rhythms in pyramidal cell excitability, these changes may not influence overall excitatory drive to PV cells in the PFC.

**Figure 11.**
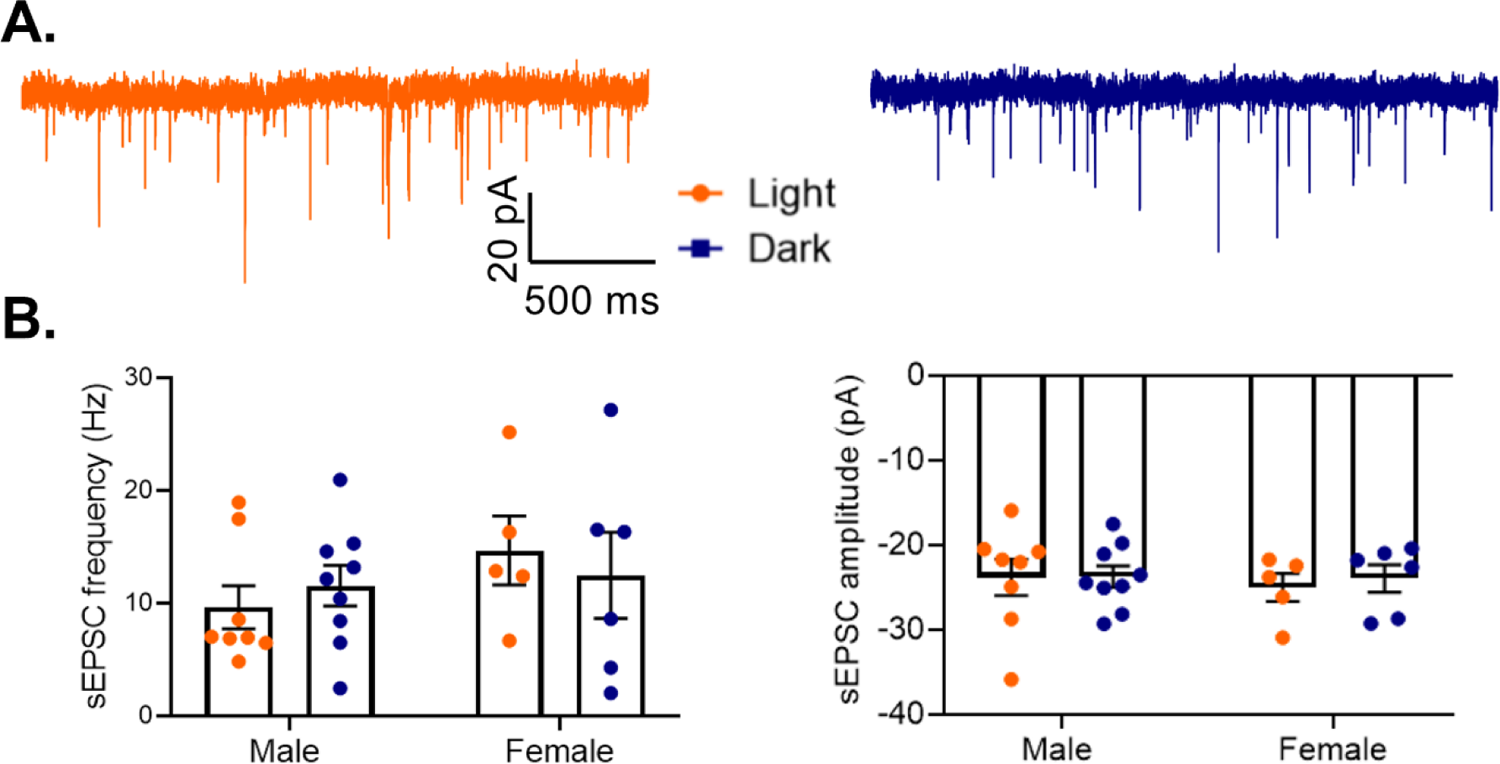
No diurnal variation in excitatory drive to PV cells (A) Representative traces of spontaneous excitatory postsynaptic currents (sEPSCs) onto PV cells in the light (left) or dark (right) phase. (B) sEPSCs onto PV cells were measured in the mouse PFC. There is no effect of phase on either sEPSC frequency (Main effect of phase: F(1,24)=0.003, p=0.96) (left) or sEPSC amplitude (Main effect of phase: F(1,24)=0.10; p=0.76)(right). n= 5-9 cells/group; 3-6 animals/group. PV=parvalbumin cells, sEPSC=spontaneous excitatory post synaptic current

### Environmental circadian desynchronization (ECD) abolishes electrophysiological rhythms in PV cells

Previous work has shown that diurnal rhythms in pyramidal cell electrophysiology are eliminated by ECD (Roberts and Karatsoreos, 2023). As we found that both action potential amplitude and spike frequency adaption show diurnal variation in PV cells from females, we utilized the same ECD paradigm to determine if these changes are vulnerable to the desynchronization of circadian rhythms. In this protocol, animals were housed under a 10:10 L:D cycle, resulting in a daily phase advance of 4 hours. We find that ECD is sufficient to abolish diurnal rhythms in both action potential amplitude (t(19)=1.08; p=0.29) and spike frequency adaptation (t(19)=0.45, p=0.66) (Fig. 12A-C).

**Figure 12.**
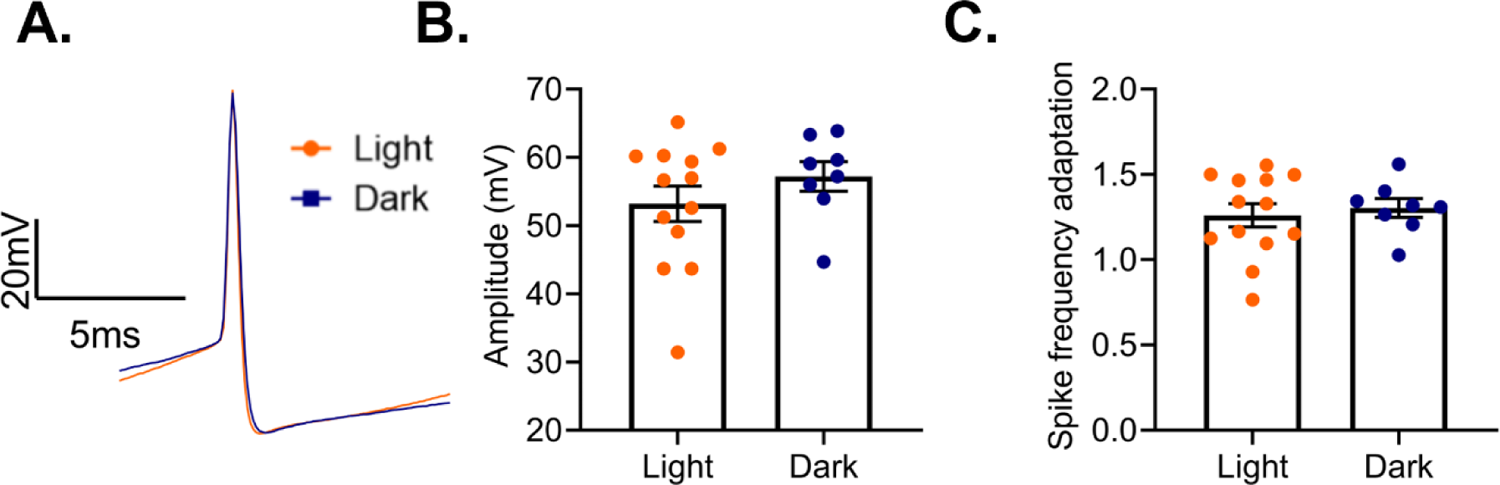
Environmental circadian desychronization eliminates rhythms in PV cell electrophysiology (A) Representative traces showing loss of diurnal variation in action potential amplitude in PV cells from female mice after environmental circadian desynchronization (ECD). (B) After ECD, there was no difference in action potential amplitude between cells recorded in the light phase or the dark phase (t(19)=1.08; p=0.29). (C) ECD resulted in a loss of diurnal variation in PV cell spike frequency adaptation (t(19)=0.45, p=0.66). n=8-13 cells/group; 3-6 animals/group.

## Discussion

In this study, we analyzed the cell-type specific diurnal rhythmicity of mRNA transcripts among two cell types that are important for PFC function and have been heavily implicated in psychiatric disease (Beasley and Reynolds, 1997; Sohal et al., 2009; Yizhar et al., 2011; Chung et al., 2016; Hoftman et al., 2017). We find that, in the PFC, the core molecular clock is synchronized between excitatory (pyramidal) and inhibitory (PV) neurons, suggesting that previously described rhythms in the E/I balance (Chellappa et al., 2016; Bridi et al., 2020) are likely not the result of these cells cycling opposing circadian phases. The synchronization of rhythms may contribute to complex behaviors that rely on these cells working together as a cohesive circuit. Indeed, previous studies found that altered rhythms in core clock genes in the PFC are accompanied by changes in mood-related behaviors, whereas other studies found that circadian desynchrony leads to deficits in cognitive flexibility (Karatsoreos et al., 2011; Otsuka et al., 2020). Another study demonstrated that disrupting rhythms through a pyramidal cell-type specific deletion of the core clock gene *Bmal1* was sufficient to impair performance in learning and memory tasks (Price et al., 2016). Together, these studies highlight the importance of rhythms on PFC-mediated functions.

Similar to studies in the SCN (Wen et al., 2020), the proportion and identity of rhythmic transcripts differs by cell type in the PFC. We find that in pyramidal cells, a high percentage of transcripts display rhythms in their expression (∼35%) and that ∼71% of all identified rhythmic transcripts in this study are rhythmic exclusively in pyramidal cells. Intriguingly, a study has shown that there is a pathway independent of the SCN that links the retina to the PFC. In this pathway, intrinsically photosensitive retinal ganglion cells synapse onto cells in the perihabenular region, which in turn send projections to the PFC (Fernandez et al., 2018). While it is not clear which cell type receives this input, our data would suggest that pyramidal cells may be directly targeted given their high level of rhythmic gene expression. As the animals used for RNA sequencing were housed under a 12:12 L:D cycle, the effect of light and the anatomy of these projections will need to be analyzed in future studies.

Consistent with studies showing that genes associated with the cell cycle are under transcriptional control of the molecular clock (Miller et al., 2007), we find that many rhythmic transcripts, particularly in pyramidal cells, are associated with cancer related pathways. However, many of these transcripts are also important for broader neuronal function. For example, studies have shown that transcripts within these pathways, such as *mTOR* and those belonging to the *Wnt* family, play an important role in synaptic plasticity (Panayiotis Tsokas et al., 2007; Hoeffer and Klann, 2010; Tabatadze et al., 2014; Narvaes and Furini, 2022).

Interestingly, studies have found that mTOR signaling is altered in the PFC of individuals with major depressive disorder and is hypothesized to underlie the rapid antidepressant effects of ketamine (Li et al., 2010; Jernigan et al., 2011). Therefore, rhythms in these transcripts may have implications for diseases beyond cancer.

In females, many of the rhythmic transcripts in PV cells belong to pathways associated with synaptic transmission. As complex behaviors rely on communication across circuits, rhythms in transcripts associated with neurotransmission may have wide ranging effects on behavior. Indeed, studies have found a time of day effect on the discrimination ratio in a novel object recognition task, which is dependent upon interaction between the hippocampus and the PFC (Cong Wang et al., 2021; Liu et al., 2022). Interestingly, rhythmic transcripts in females are particularly enriched for processes such as synaptic long-term potentiation/depression during the inactive phase. This corresponds with the time in which the perineuronal net (PNN), a specialized form of the extracellular matrix that forms around PV cells and restricts plasticity, is decreased (Carulli et al., 2010; Wang and Fawcett, 2012; Pantazopoulos et al., 2020; Harkness et al., 2021). Therefore, reduction in the PNN during the light phase may allow for increased plasticity and may play a role in the synaptic remodeling that occurs during sleep (Tononi and Cirelli, 2006; Vyazovskiy et al., 2008; Choudhury et al., 2020).

We find that rhythmic transcripts in PV cells from males are highly enriched in processes associated with mitochondrial function and oxidative phosphorylation. This suggests that there might be a unique cell-type and sex-specific vulnerability of PV cells in males to the production of reactive oxygen species in response to environmental changes in the sleep/wake cycle.

Interestingly, studies in mice have suggested that PV cells are particularly vulnerable to oxidative stress, and it has been proposed that oxidative stress induced damage to PV cells may be a key component of schizophrenia pathophysiology (Behrens and Sejnowski, 2009; Cabungcal et al., 2013; Steullet et al., 2017). Work using human postmortem tissue, primarily from male subjects, has shown that there are changes in the transcriptome of PV cells in subjects with schizophrenia, with genes associated with oxidative phosphorylation and mitochondrial dysfunction being among the most differentially expressed (Enwright Iii et al., 2018). Moreover, studies have shown that individuals with schizophrenia show vast circadian reprograming across multiple brain regions and that differential expression of mitochondrial related transcripts in schizophrenia can be explained by altered patterns of rhythmicity (Seney et al., 2019; Ketchesin et al., 2023). Therefore, the enrichment of rhythmic transcripts associated with mitochondrial function in PV cells in males suggests a potential mechanism whereby altered molecular rhythms in schizophrenia may impact normal diurnal rhythms in oxidative phosphorylation in PV cells, leaving these cells vulnerable to oxidative stress.

Strikingly, there is more overlap in rhythmicity between PV and pyramidal cells from female animals than within PV cells across sexes. This suggests that there is a sex-specific role of rhythmic transcripts that is conserved across cell types. Previous studies have found that the *Clock* and *Per2* promoters contain estrogen response elements and that their expression is modulated through the estrogen receptor ESR1 (Hatcher et al., 2020; Alvord et al., 2022). This suggests that interactions between the molecular clock and estrogen signaling may coordinate rhythms, contributing to the overlap in rhythmic transcripts across cell types in females. Notably, we find that *ESR1* is not differentially expressed between sexes in pyramidal cells (q=0.22, FC=1.43). Conversely, in PV cells, *ESR1* expression differs by sex (q=0.00, FC=1.72), perhaps allowing circulating estrogens to have a greater effect on rhythms and contributing to the broad sex differences in rhythmicity that we find in PV cells.

Nevertheless, across cell types and sexes, there are 116 transcripts that are rhythmic in all groups (Table 1-15). These conserved rhythmic transcripts show enrichment for pathways associated with circadian rhythm signaling and include many of the top rhythmic transcripts. Other conserved transcripts are associated with basic biological processes such as translation and protein folding/degradation (e.g. *Eif4ebp1, Hsf1*, *Hspa1b, Hspa5, Usp2, Usp43*) and MAP kinase mediated signal transduction (e.g. *Map2k3, Mapk11*), emphasizing the importance of rhythms in these processes across cell types.

Electrophysiological recordings serve as a tool to study the functional output of rhythms in gene expression. A previous study examining the effect of time of day on pyramidal cell electrophysiology in the PFC found that there are sex-specific effects on resting membrane potential, action potential frequency, threshold, half-width, and membrane conductance (Roberts and Karatsoreos, 2023).Therefore, we performed complementary recordings in PV cells to determine if similar rhythms exist in PV cell intrinsic properties. Consistent with the lower proportion of rhythmic transcripts in PV cells relative to pyramidal cells, our results indicate more subtle diurnal changes in PV cell electrophysiological properties. We find no effect of phase on properties such as firing frequency, rheobase, input resistance, half-width, or afterhyperpolarization amplitude. However, in females, we find that both action potential amplitude and spike frequency adaptation show significant diurnal variation, which is eliminated by ECD. This protocol has previously been shown to lead to a loss of rhythms in pyramidal cell electrophysiology as well to changes neuronal morphology in the PFC and deficits in cognitive flexibility (Karatsoreos et al., 2011; Roberts and Karatsoreos, 2023). Combined, this work provides insight into how circadian desynchrony, such as during shift work, may contribute to impairments in cognitive function through disruption of rhythms in neuronal physiology. Notably, while many of the observed time of day effects in pyramidal cells are present only in males (Roberts and Karatsoreos, 2023), diurnal changes in PV cell electrophysiology are driven by females, suggesting that the effects of circadian disruption may differ by sex.

Previous studies have found that diurnal variation in the frequency of excitatory and inhibitory transmission occurs in opposing phases in PV and pyramidal cells in the cortex (Bridi et al., 2020; Zong et al., 2023). However, in this study, we find that there is no diurnal change in the excitatory drive to PV cells. As the diurnal variation in the E/I ratio onto pyramidal cells is driven by local, lateral connections (Bridi et al., 2020), future studies should be done to parse the role of different PV cell inputs on the E/I balance, both from the PFC and other regions. Moreover, our finding of no diurnal change in the amplitude or frequency of sEPSCs does not preclude more subtle changes, such as changes in receptor kinetics. Indeed, we find that there are sex-specific rhythms in the expression of multiple AMPA receptor subunits in PV cells, whereby *Gria3* and *Gria4* are rhythmic in females, while *Gria1* is rhythmic in PV cells from males. In conclusion, in this study, we show that there are both cell-type and sex-specific differences in rhythmic gene expression and electrophysiological properties in the mouse PFC. We propose that these represent unique roles of rhythms in cell-type and sex-specific processes. Moreover, the unique biological processes associated with rhythmic transcripts may provide insight into the selective vulnerabilities of different cell types to circadian disruption, which can be further investigated through cell-type specific manipulation of the molecular clock.

## Extended Data

**Table 1-1:** Rhythmic transcripts (p<0.05) in pyramidal cells with sexes combined

**Table 1-2:** Rhythmic transcripts (p<0.05) in PV cells with sexes combined

**Table 1-3:** Rhythmic transcripts (p<0.05) from pyramidal cells in females

**Table 1-4:** Rhythmic transcripts (p<0.05) from pyramidal cells in males

**Table 1-5:** Uniquely rhythmic transcripts from pyramidal cells in males (vs. females)

**Table 1-6:** Uniquely rhythmic transcripts from pyramidal cells in females (vs. males)

**Table 1-7:** Rhythmic transcripts (p<0.05) from PV cells in females

**Table 1-8:** Rhythmic transcripts (p<0.05) from PV cells in males

**Table 1-9:** Uniquely rhythmic transcripts from PV cells in males (vs. females)

**Table 1-10:** Uniquely rhythmic transcripts from PV cells in females (vs. males)

**Table 1-11:** Uniquely rhythmic transcripts from pyramidal cells in females (vs. PV cells)

**Table 1-12:** Uniquely rhythmic transcripts from PV cells in females (vs. pyramidal cells)

**Table 1-13:** Uniquely rhythmic transcripts from pyramidal cells in males (vs. PV cells)

**Table 1-14:** Uniquely rhythmic transcripts from PV cells in males (vs. pyramidal cells)

**Table 1-15:** Transcripts that are rhythmic (p<0.05) in both pyramidal and PV cells in both sexes

## Acknowledgements

This work was funded by MH106460, NS127064, DA039865, DA046346, MH111601 and the Wood Next Foundation to CAM. We would like to thank the University of Pittsburgh Health Sciences Sequencing Core and UPMC Genome Center for assistance with library preparation and RNA-sequencing. We thank Yianni Migias for animal care and husbandry. The authors declare no competing financial interests.

